# Probing the association between resting state brain network dynamics and psychological resilience

**DOI:** 10.1101/2021.07.20.452941

**Authors:** Dominik Kraft, Christian J. Fiebach

**Affiliations:** Department of Psychology, Goethe University Frankfurt, Germany; Brain Imaging Center, Goethe University Frankfurt, Frankfurt am Main, Germany

**Keywords:** resting state, time varying connectivity, multilayer modularity, psychological resilience, network reconfigurations, node flexibility, node promiscuity, node degree

## Abstract

This study aimed at replicating a previously reported negative correlation between node flexibility and psychological resilience, i.e., the ability to retain mental health in the face of stress and adversity. To this end, we used multiband resting-state BOLD fMRI (TR = .675 sec) from 52 participants who had filled out three psychological questionnaires assessing resilience. Time-resolved functional connectivity was calculated by performing a sliding window approach on averaged time series parcellated according to different established atlases. Multilayer modularity detection was performed to track network reconfigurations over time and node flexibility was calculated as the number of times a node changes community assignment. In addition, node promiscuity (the fraction of communities a node participates in) and node degree (as proxy for time-varying connectivity) were calculated to extend previous work. We found no substantial correlations between resilience and node flexibility. We observed a small number of correlations between the two other brain measures and resilience scores, that were however very inconsistently distributed across brain measures, differences in temporal sampling, and parcellation schemes. This heterogeneity calls into question the existence of previously postulated associations between resilience and brain network flexibility and highlights how results may be influenced by specific analysis choices.

**Author Summary:** We tested the replicability and generalizability of a previously proposed negative association between dynamic brain network reconfigurations derived from multilayer modularity detection (node flexibility) and psychological resilience. Using multiband resting-state BOLD fMRI data and exploring several parcellation schemes, sliding window approaches, and temporal resolutions of the data, we could not replicate previously reported findings regarding the association between node flexibility and resilience. By extending this work to other measures of brain dynamics (node promiscuity, degree) we observe a rather inconsistent pattern of correlations with resilience, that strongly varies across analysis choices. We conclude that further research is needed to understand the network neuroscience basis of mental health and discuss several reasons that may account for the variability in results.

## Introduction

Abundant literature in human clinical neuroscience has established a link between changes in intrinsic functional connectivity of large-scale brain networks and psychological disorders such as depression (Mulders et al., 2015) or schizophrenia (Dong et al., 2018; see also Menon, 2011 for a general overview). As a consequence, brain bases of preserving mental health in the face of stress and adversity (resilience) have also become a focus of interest (e.g., Southwick & Charney, 2012). Two recent studies (Long et al., 2019; Paban et al., 2019) have reported associations between psychological resilience and brain network dynamics using multilayer modularity, a relatively new tool from the evolving field of network neuroscience that integrates spatial and temporal information (Muldoon & Bassett, 2016). In both studies, resilience was assessed with a frequently-used questionnaire, the Connor-Davidson Resilience Scale (CD-RISC; Connor & Davidson, 2003), and network dynamics under task-free (resting-state) conditions were examined by detecting time-evolving patterns of non-overlapping and coherent modules and by quantifying the frequency with which brain nodes switched between modules (*node flexibility;* Bassett et al., 2011).

The link between resilience and brain network dynamics is motivated (a) by an assumed relationship between neuronal and cognitive flexibility (e.g., Braun et al., 2015; Chen et al., 2016) and (b) given that resilience has been associated with higher cognitive flexibility, both in theoretical models (Southwick & Charney, 2012) and in empirical work (e.g., Genet & Siemer, 2011). While this would predict greater resilience in more (cognitively or neuronally) flexible persons, it has also been argued that cognitive flexibility might not be universally adaptive and that resilience may depend on an interplay between flexibility and stability (Parsons et al., 2016). This proposal receives support from observations of changes along flexibility-stability dimensions in psychiatric conditions, both behaviorally (e.g., rigid behaviors like rumination in depression; Nolen-Hoeksema et al., 2008) and neuronally (e.g., increased network flexibility, variability in patients with autism or schizophrenia; Gifford et al., 2020; Harlalka et al., 2019). These and similar results suggest that the extreme tails of the ‘flexibility-stability’ distribution (Kashdan & Rottenberg, 2010) may indeed be related to psychopathology and that, accordingly, adaptive behavior in the ‘normal’ (i.e., unaffected) range depends on a balance between flexibility and stability. Whereas the above-cited work by Long et al. (2019) and Paban et al. (2019) took a primarily neuroscientific perspective as a starting point, the psychological perspective offered here is not fully consistent with their empirical results. However, it may offer a valuable conceptual framework for a neuro-cognitive model of resilience and therefore calls for further empirical research to clarify the current inconsistencies.

Paban et al. (2019) measured task-free EEG and conducted network analyses in source space. Negative correlations between node flexibility and resilience were observed in the alpha, beta, and delta frequency bands, including superior parietal cortex, medial orbitofrontal cortex, and cuneus. A subsequent resting state functional MRI (rs-fMRI) study (Long et al., 2019) also reports lower node flexibility in more resilient persons, primarily in visual cortices and the left medial-orbital superior frontal gyrus. Except for a partial overlap in visual regions (lingual gyrus), localization results differed between studies and Long et al. (2019) did not replicate correlations in ‘higher order’ (superior parietal, inferior frontal) areas reported by Paban et al. (2019). Whereas some of these inconsistencies may result from inherent differences between methods (e.g., differences in temporal resolution between fMRI and EEG), others may reflect specific methodological choices by the authors. For example, Long et al. (2019) studied BOLD ‘dynamics’ using only 12 non-overlapping time windows (each of 20 sample points length, derived from 250 measurements of TR = 2 sec).

Even though gold standards for analyses of dynamic multilayer modularity are yet to be established, recent studies suggest that more data are required for sensitive and reliable estimation of network dynamics from BOLD fMRI (Hindriks et al., 2016; Yang et al., 2021). To ameliorate such methodological shortcomings and to further our understanding of the relationship between resilience and brain network dynamics, we here replicate and extend these results using temporally highly resolved fMRI (multiband/MB sequence; MB factor 4; TR = .675 sec) from 52 healthy young adults who completed three resilience questionnaires. We first replicated Long et al.’s (2019) analysis pipeline as closely as possible, by down-sampling data to a TR of 2.025 sec and using the same analysis parameters. Following this, we explore effects of specific analysis choices (like different windowing schemes) on network flexibility. Lastly, we repeat correlation analyses with optimized denoising and the full temporal resolution of the MB data, resulting in a timeseries of 753 overlapping windows (Figure 1). Node flexibility was calculated as the number of times a node changes its community assignment between windows, divided by the total possible number of changes (Bassett et al., 2011). Given the results of Long et al. (2019), we expected that resilience and node flexibility should be negatively correlated.

**Figure 1.**
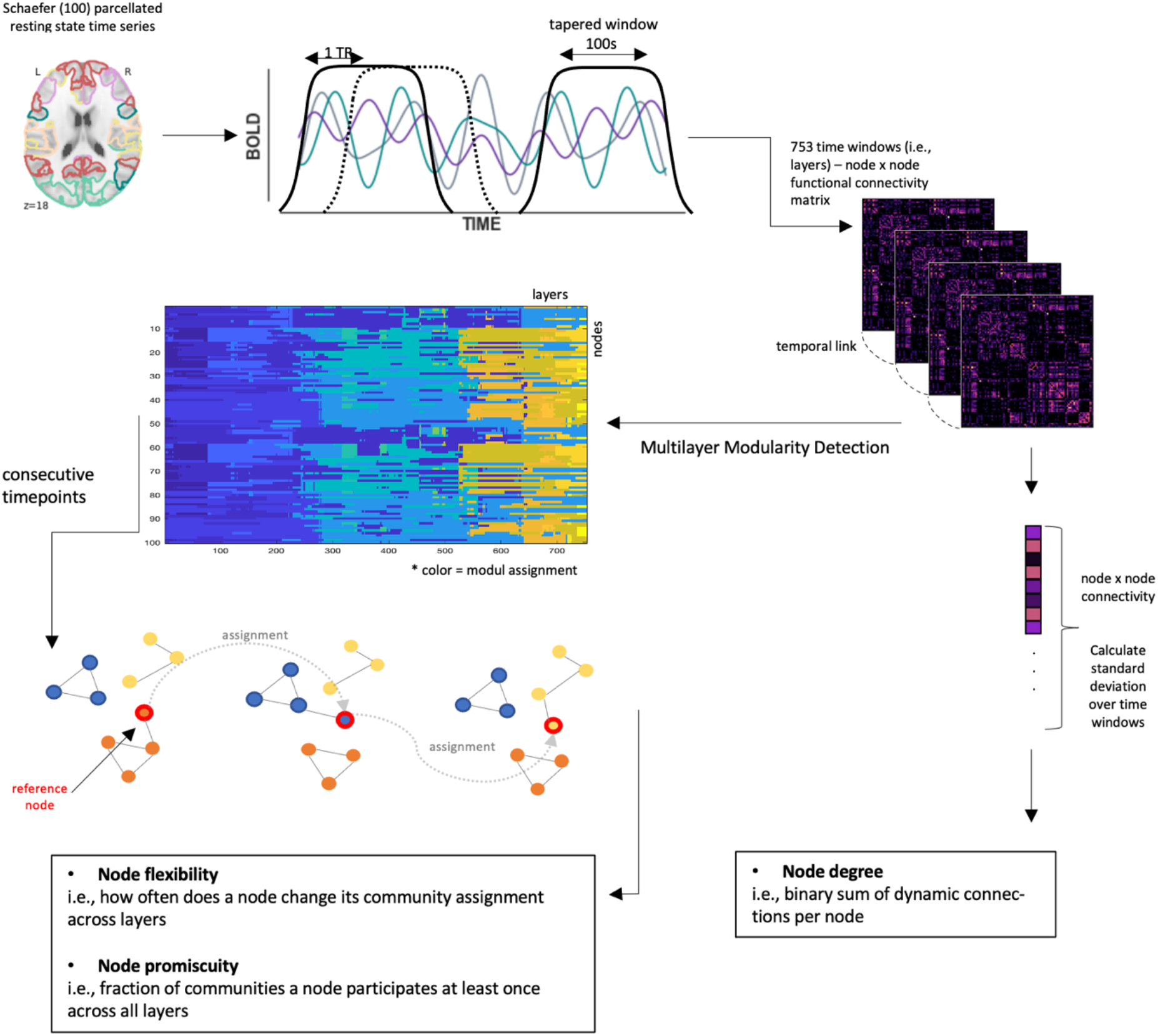
Schematic illustration of the workflow, here visualized for analyses with original multiband data (TR = .675 sec), the Schaefer100 atlas, and overlapping sliding windows of size 100 sec and offset 1 TR (see *Methods* for details). Procedures for other parameter choices are analogous. Starting with the mean resting state time series, 753 functional connectivity matrices (layers) representing inter-correlations between the 100 different nodes were calculated via a sliding-window approach; multilayer modularity detection was performed on ordinal layers. Network measures were calculated as described in the figure; see *Methods* section for further details

To extend previous work, we also assessed node promiscuity and node degree, and their relation to resilience. Node promiscuity, the fraction of communities a node participates in at least once, and node flexibility are complementary measures that inform us whether brain dynamics per se (i.e., assessed via node flexibility) or the diversity of brain systems with which a node interacts (assessed via its promiscuity) may be associated with resilience. Node degree, in turn, is a proxy for dynamic connectivity that does not rely on modularity detection algorithms (and is thus invariant to potential algorithmic idiosyncrasy or parameter choices). However, previous work demonstrates that dynamic connectivity is related to various cognitive and behavioral traits, as well as clinical conditions like post-traumatic stress disorder (PTSD, often used as proxy for studies of resilience; Bolsinger et al., 2018; Jin et al., 2017; Lurie et al., 2020), which makes it a candidate marker for analyzing the neurobiology of resilience. To the best of our knowledge, relationships between resilience and promiscuity as well as degree have so far not been studied. Accordingly, we here aim at a more complete characterization of putative relationships between brain network dynamics and resilience.

## Results

### Behavioral Results: Subjective Resilience Ratings

Descriptive statistics for resilience scales are listed in Table 1. Psychometric properties were satisfactory, with internal consistencies (Cronbach’s alpha) between .7 and .85 (Table 1). Resilience scores in our sample were comparable to the respective original publications for the German versions (CD-RISC: 30.6, Sarubin et al., 2015; RS-13: 70, Leppert et al., 2008; BRS: 3.58 and 3.37, Chmitorz et al., 2018). Resilience ratings were significantly intercorrelated: CD-RISC – RS, *r =* .60; CD-RISC – BRS, *r =* .61; BRS – RS, *r =* .45, all *p* < .001 (FDR-corrected for multiple comparisons).

**Table 1.** Descriptive statistics of resilience questionnaires (N = 52)

|  | **CDRISC** | **RS** | **BRS** |
| --- | --- | --- | --- |
| *M* (*SD*) | 28.88 (4.73) | 71.37 (10.06) | 3.72 (.73) |
| min – max | 18 - 38 | 21 - 87 | 1.83 – 5.00 |
| $\alpha$ (95% CI) | .70 (.56- .81) | .85 (.79- .91) | .82 (.73-.89) |

Note: M = mean, SD = standard deviation, min = minimum, max = maximum, $\alpha$ = Cronbach's alpha, CI = confidence interval.

### Replication of Long et al. (2019)

We replicated the analysis pipeline of Long et al. (2019) as closely as possible, involving down-sampling of multiband data by a factor of 3 to a virtual TR of 2.025 seconds, use of the two parcellation schemes used by Long et al. (2019), i.e., AAL90 and Power264 atlases, and the same denoising strategy (with 26 parameters) as used in that study. Age, gender, and framewise displacement were included as covariates of no interest (see Methods). Given that recent evidence implies that functionally derived parcellations outperform anatomical atlases like AAL (Dadi et al., 2019), we also conducted all analyses with a further atlas, the functionally defined parcellation scheme of Schaefer (100 nodes) which approximately matches the number of nodes in the AAL. Following Long et al. (2019), the down-sampled timeseries was segmented into non-overlapping windows of 20 TRs length, resulting in 15 time windows (as opposed to 12 in the original study). We did not replicate the correlation between global flexibility and CD-RISC reported by Long et al. (2019; all *p* >= .33), but observe a borderline significant correlation between global flexibility (AAL atlas) and the BRS resilience score (*r* = −.33, *p* = .05). Across the three parcellation schemes and resilience questionnaires, we did not observe any further significant effects, neither at global, subnetwork, or node level (Table 2).

**Table 2:** Correlations between global flexibility and resilience scales according to the pipeline used in Long et al. (2019)

| atlas | resilience scales |  |  |
| --- | --- | --- | --- |
|  | CD-RISC | BRS | RS |
| AAL 90 | $r = -.14, p = .33$ | $r = -.33, p = .05$ | $r = -.20, p = .23$ |
| Power 264 | $r = .06, p = .87$ | $r = -.02, p = .87$ | $r = -.03, p = .87$ |
| Schaefer 100 | $r = -.01, p = .94$ | $r = -.12, p = .58$ | $r = -.16, p = .58$ |

### Effects of overlapping vs. non overlapping time windows

Segmenting the timeseries into non-overlapping windows results in a low number of sample points, generally not considered sufficient for analyzing network dynamics. Many studies today segment BOLD timeseries into overlapping windows, as this allows for an estimation of network dynamics with greater sensitivity and reliability (Hindriks et al., 2016; Lurie et al., 2020; Yang et al., 2021). When analyzing our down-sampled data with a high number of overlapping windows (i.e., 251; see Methods), nodal flexibility was scaled by a factor of ∼1/10; cp. y axis scaling of Figures 2A and 2B), as expected given the higher overlap between consecutive windows. However, high spatial similarity was preserved between node flexibility derived from down-sampled data with overlapping vs. non-overlapping windows (*r* = .90, *p* < .0001; Figure 2C,D), indicating that there are no qualitative differences between windowing schemes. We thus use overlapping windows for all further analyses.

**Figure 2.**
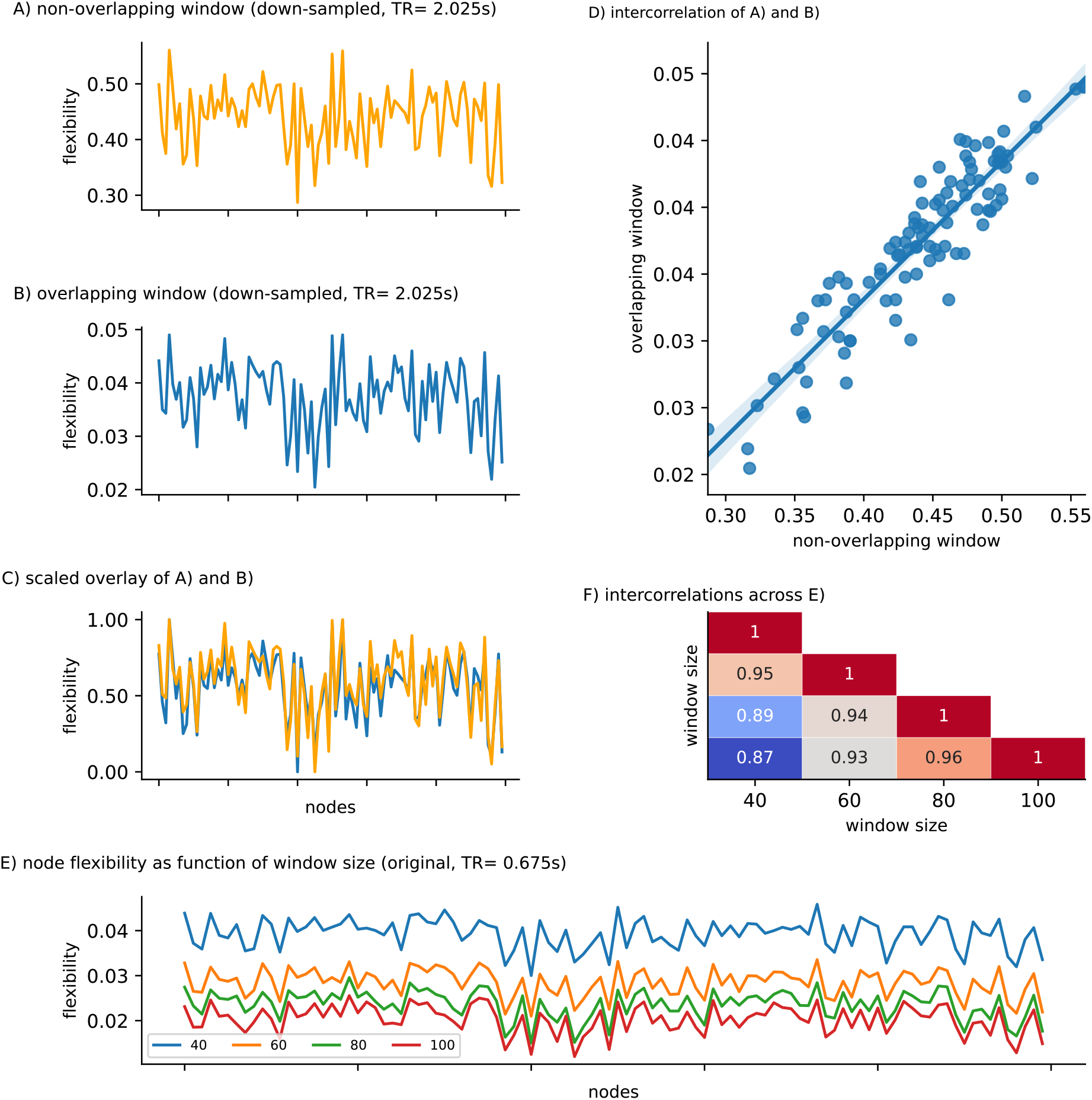
Influence of methodological differences in dynamic network analysis on node flexibility. Node flexibility for the down-sampled data (TR = 2.025 sec) with (A) non-overlapping (window size = 20TR) vs (B) overlapping windows (window size = 50TR). Overlaying rescaled variants of both arrays indicates (C) only minor differences between node flexibility values derived from different window schemes and (D) high spatial similarity, i.e., a high correlation coefficient, between (A) and (B). (E) Node flexibility for the original data (multiband, TR = .675 sec), across nodes (x axis) as a function of window size (40 to 100 sec length) using overlapping windows (F) The distribution of flexibility values across nodes shows high spatial similarity (all correlations *p* < .0001). For the analyses shown in (E) and (F), flexibility values were derived after repeating the modularity detection algorithm analyses 10 times and choosing the run yielding the highest modularity value Q (see *Methods* for details).

### Dynamic network measures in original vs. down-sampled functional connectivity data

Analyses of network dynamics may be limited by lower temporal resolutions (which was here simulated by downs-sampling to allow for a direct replication of the results by Long et al., 2019). As compared to the original data (TR = .675 sec), down-sampling shifted node flexibility (calculated using over-lapping windows; see previous section) towards higher values. However, high spatial similarity (i.e., correlation) exists between the distribution of nodal flexibility in original vs. down-sampled data (*r* = .95, *p* < .0001). As outlined above, we will in the following also investigate node promiscuity and degree as further measures of brain network flexibility. We thus also explore how these measures are affected by down-sampling of the BOLD data: As for flexibility, down-sampling led to higher promiscuity values, but a high spatial similarity was preserved (*r* = .93, *p* < .0001). We observed a significantly lower nodal degree in the down-sampled data (0.06 – 0.61% significant connections; *M =* 0.20) compared to the original data (0.12 – 0.75%; *M =* 0.33), *t(99)* = 13.93, *p* < .0001, which is consistent with a recent report by Pedersen et al. (2018) that fewer data result in lower nodal degree values.

### Correlation between dynamic functional connectivity measures and resilience

Given that no previous data exist to derive specific hypotheses concerning the relationship between resilience and node promiscuity as well as degree, all subsequently reported associations were tested two-sided using Spearman correlations. Correlations were calculated for all three parcellations, for original (TR = .675 sec) and down-sampled (TR = 2.025) data, using an optimized denoising strategy (36 parameters; see Methods), overlapping time windows, and the same covariates as during the replication attempt reported above.

In the original data, we observed two significant positive correlations at the nodal level of the Power264 atlas between flexibility of a cingulo-opercular node (#54) and the CD-RISC (*r* = .52, *p* = .02), and between promiscuity of a visual node (#165) and the BRS resilience score (*r* = .57, *p* = .004). After down-sampling (but now exploiting higher temporal resolution due to the overlapping windowing scheme), the AAL-based analysis yielded a significant association between the promiscuity of the left pallidum (node level, AAL) and BRS resilience (*r* = −.49, *p* = .02). In the Schaefer100 analysis, significant negative correlations emerged between the degree of a visual node (left Vis_7) and the RS score (*r* = −.48, *p* = .03), as well as between global degree and BRS resilience (*r* = −.35, *p* = .03). In the Power264 atlas we observed two borderline significant results, between the BRS resilience score and the degree of the subcortical RSN (*r* = −.37, *p* = .05) and the cerebellar RSN (*r* = −.37, *p* = .05). No correlation was observed for any additional tested combination (see Table 3). Figures 3-5 visualize nodal and RSN results in an exemplary manner for the Schaefer100 parcellation.

**Table 3:**
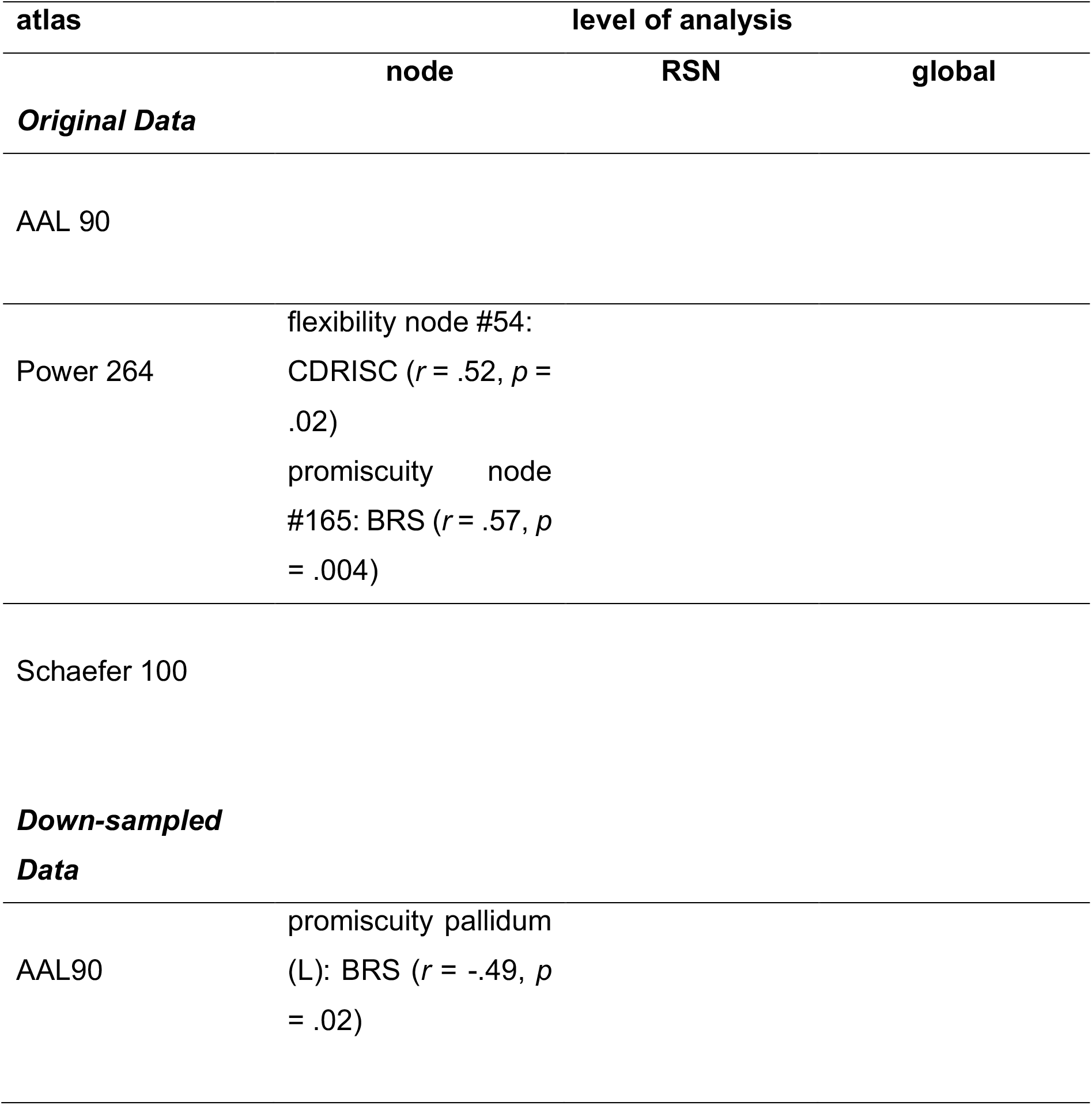
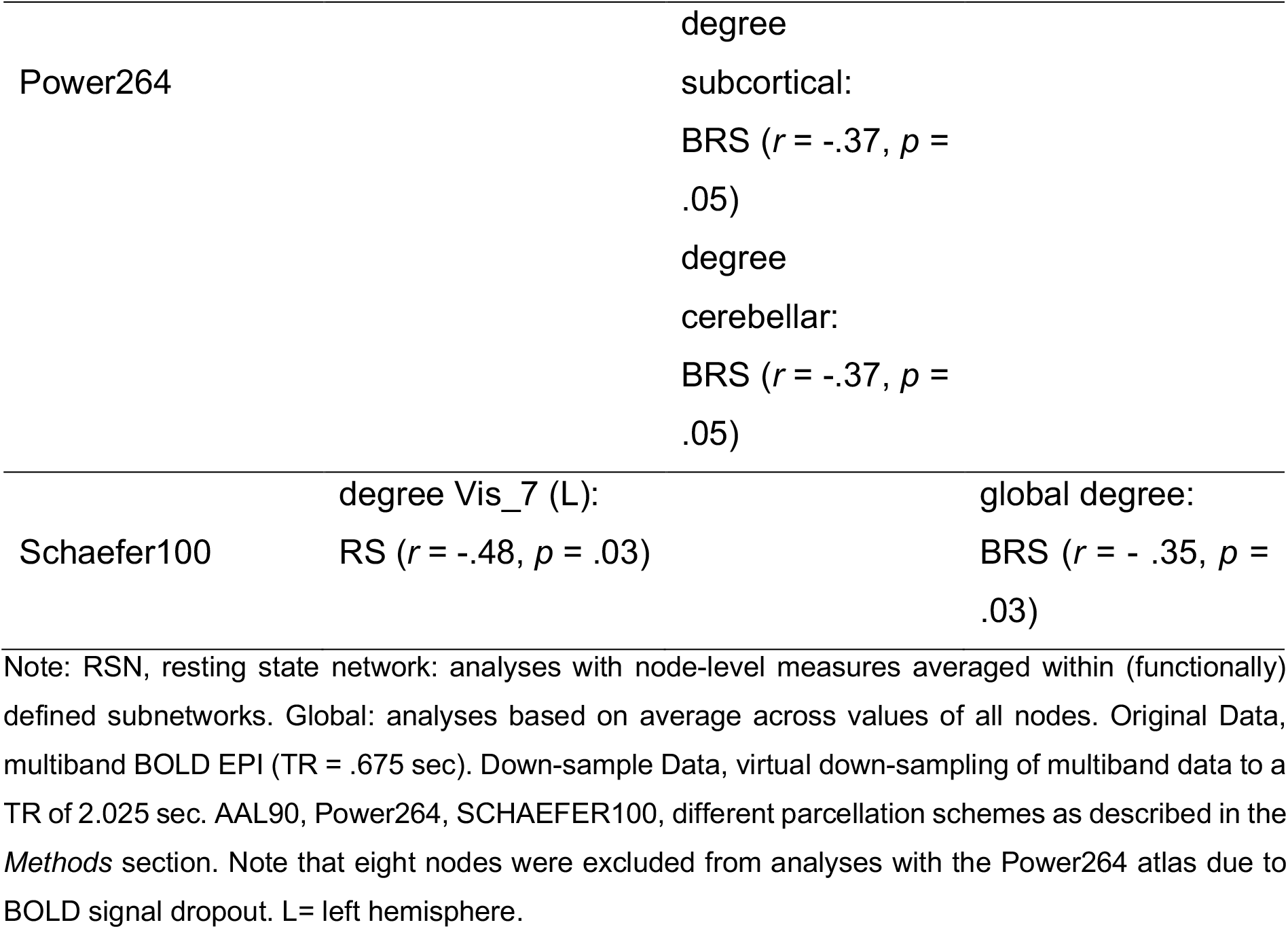
Significant correlations between measures of brain dynamics (node flexibility, node promiscuity, and node degree) and psychological resilience as measured with three different scales (CDRISC, BRS, RS; see *Methods* for details). All combinations of atlases and levels of analyses were computed, but only significant results are described in the table.

**Figure 3.**
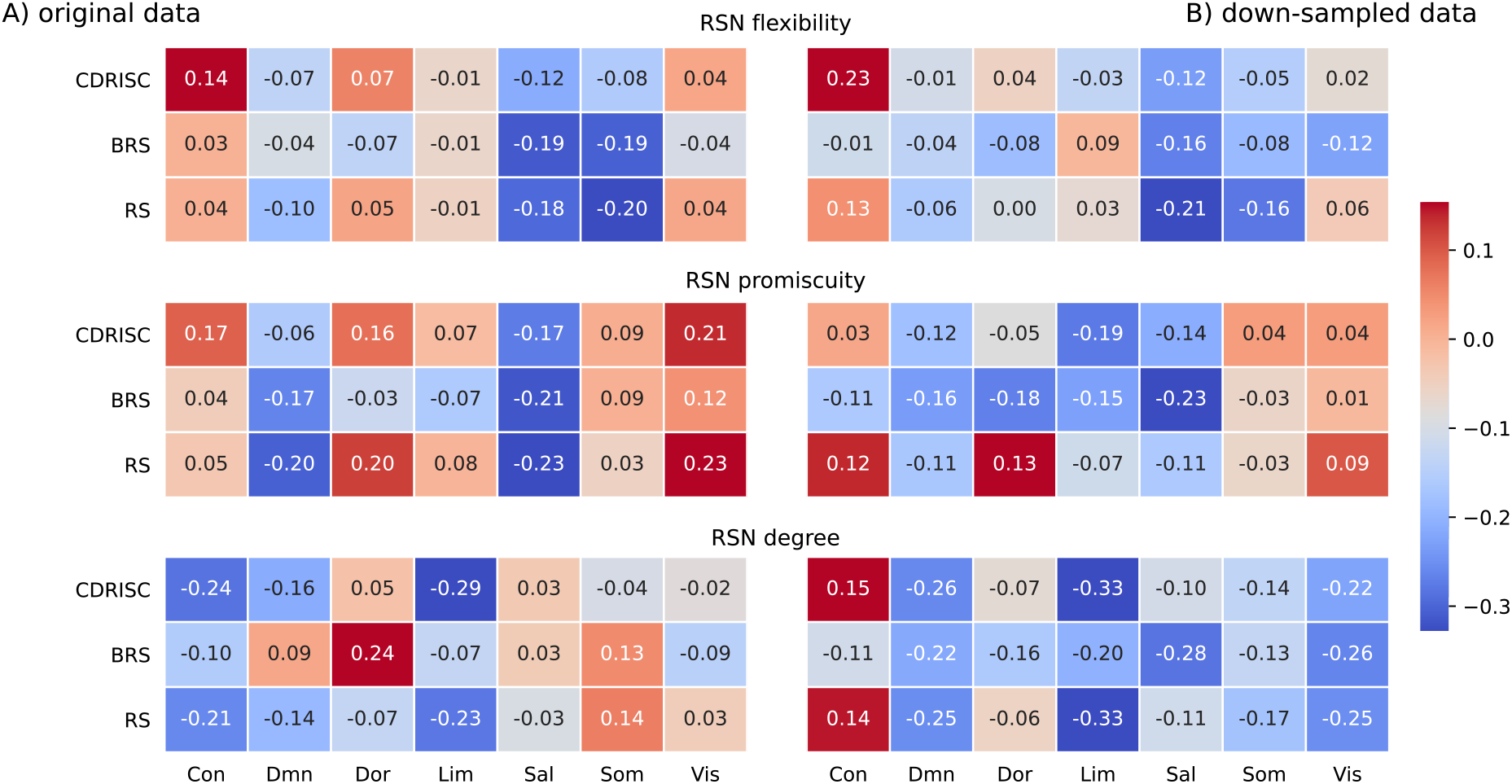
Correlations between network measures and resilience at the level of resting state networks (RSN) in the Schaefer100 atlas for the (A) original data (multiband; TR = .675 sec) and (B) down-sampled data (TR = 2.025 sec; see *Methods* for details). The number in each cell represents the respective correlation coefficient; all *p* > .12. Network labels: Con = frontoparietal-control, Sal = salience/ventral attention, Lim = limbic, Dor = dorsal attention, Som = somatomotor, DMN = default mode, Vis = visual. Resilience questionnaires: CD-RISC = Connor-Davidson Resilience Scale, BRS = Brief Resilience Scale, RS = Resilience Scale.

**Figure 4.**
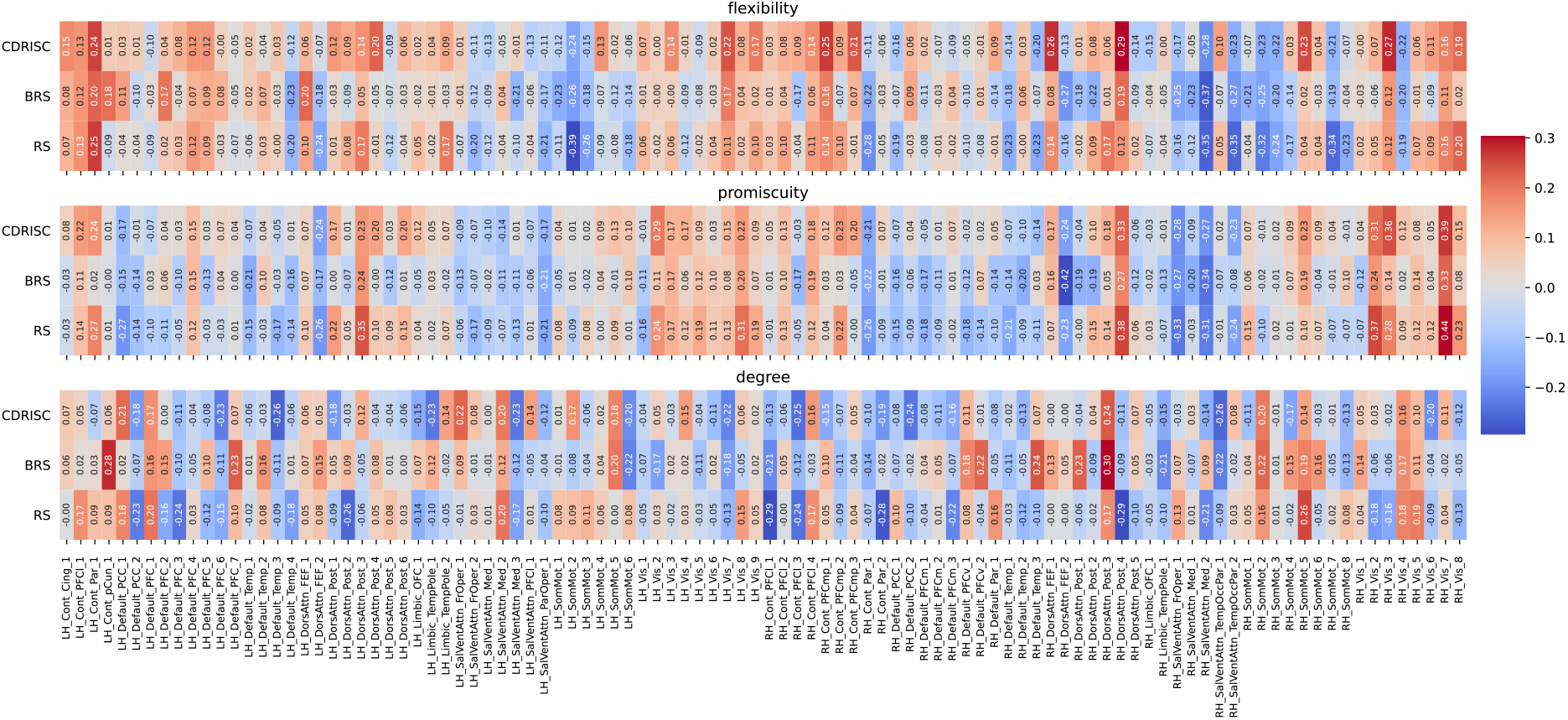
Correlations between network measures and resilience at the nodal level for the original (multiband) data (TR = .675 sec) in the Schaefer100 atlas. Node names are depicted on the x-axis ticks; for a detailed list see https://bit.ly/3yvOBwz. LH = left hemisphere, RH = right hemisphere. Figure S1 in the Supplemental Material depicts the same correlations, but without including covariates (see *Follow-up Analyses* in the *Results* section).

**Figure 5.**
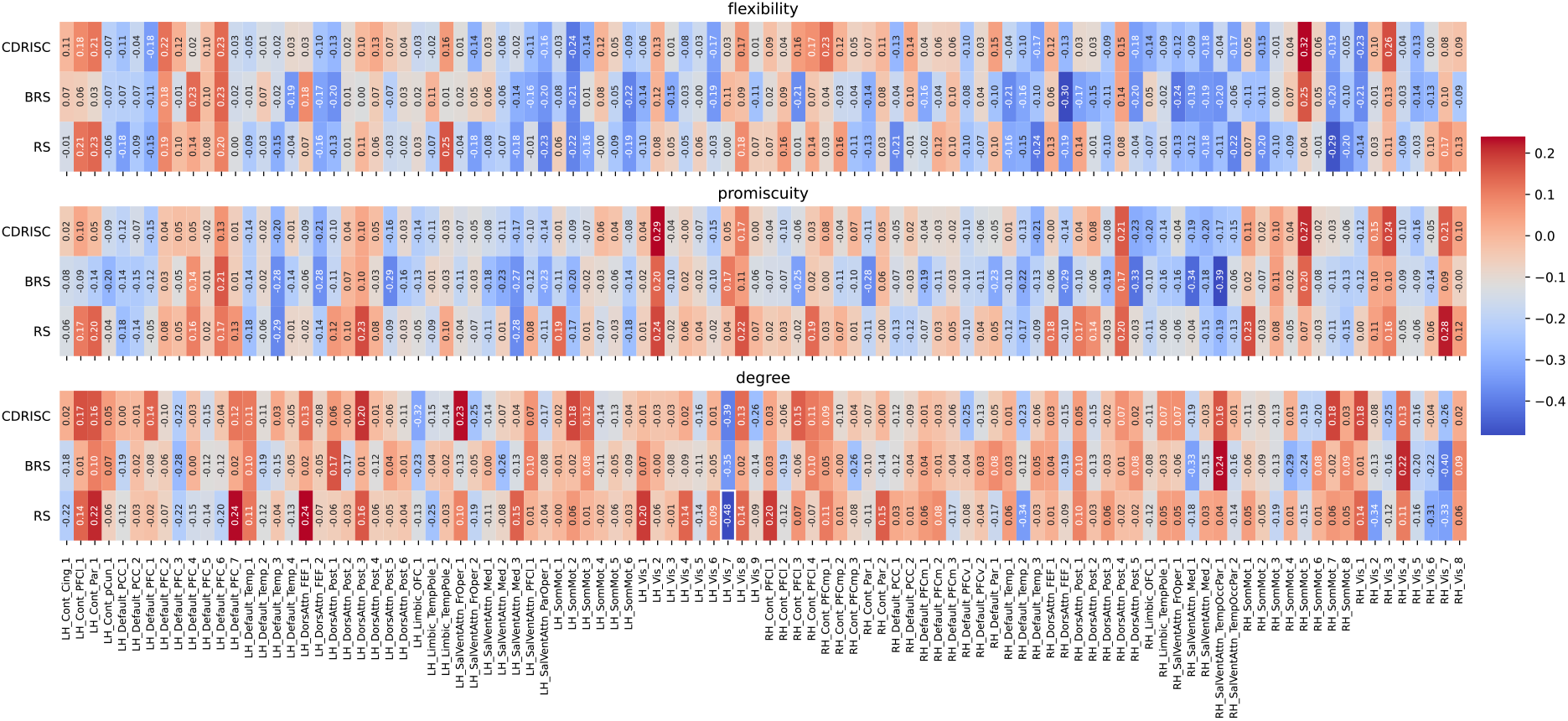
Correlations between network measures and resilience at the nodal level for the down-sampled data (TR = 2.025 sec) in the Schaefer100 atlas (all *p* > .10). Node names are depicted on the x-axis ticks; for a detailed list see https://bit.ly/3yvOBwz. LH = left hemisphere, RH = right hemisphere. The significant correlation between the degree of node Vis_7 and the RS scale is highlighted by white rectangle. Figure S2 in the Supplemental Material depicts the same correlations, but without including covariates (see *Follow-up Analyses* in the *Results* section).

### Follow-up Analyses

To further investigate putative factors that may have influenced reported results, we conducted several post-hoc analyses on data derived from the Schaefer100 parcellation.

#### Effects of covariates

Re-calculation of all node-level correlations without covariates yielded of results comparable to the above-reported findings (cf. Supplementary File 1, Figures S1 and S2). Correlations for original (MB) resolution: −.41 < *r* < .44, all *p* > .11. For down-sampled data, the above-reported significant correlation for one visual node disappeared (−.43 < *r* < .30, all *p* > .14).

#### Effects of window length

To explore how node flexibility changes as a function of window length, node flexibility was calculated for window lengths between 40 and 100 sec in steps of 20 (original/MB data). Similar to the effect of down-sampling (see above), we observed an increase in node flexibility with decreasing window size (Figure 2E), however with high spatial similarity of flexibility values across window sizes, all *r* > .87, *p* < .0001 (Figure 2F). We did not observe any significant correlations between node flexibility and resilience when varying the window size between 40 and 80 sec (all *p* > .13).

#### Effects of denoising

While repeating the correlation analyses (Schaefer100 parcellation, overlapping windows) after denoising with a 26-parameter regression model (following Long et al., 2019) as opposed to the 36-parameter model used above, no correlations with resilience were found at global (multiband: all *p* > .54; down-sampled: all *p* > .74) or nodal level (multiband: all *p* > .15; down-sampled: all *p* > .53). In the RSN analysis, significant negative correlations were found between degree of the limbic RSN and all resilience scales (multiband; CD-RISC: *r* = .-39, *p* = .03; BRS: *r* = .-44, *p* = .008; RS: *r* = .-40, *p* = .03; down-sampled: all *p* > .16).

#### Effects of spatial resolution

As the Schaefer atlas is available in multiple parcellations, it is best-suited to investigate the influence of spatial resolution. Using 200 (vs. 100) nodes, we did not observe any correlation on the global level, neither for original (all *p* > .50), nor down-sampled (all *p* > .13) data. While there were also no significant correlations on the RSN level for original data (all *p* > .29), degree of the limbic RSN was negatively correlated to both CD-RISC (*r* = −.57, *p* < .001) and RS (*r* = −.38, *p* = .04) in down-sampled data. No significant effects emerged at nodal level (original: all *p* > .36; down-sampled: all *p* > .10).

#### Effects of motion

For the 26-parameter denoised data, we observed a significant correlation between global flexibility and mean FD (*r* = .38, *p* = .02, FDR-corrected) derived from the original (multiband) data, whereas no other brain measure was correlated with FD for original or down-sampled data (all *p* > .11) and no correlations emerged after optimized (36-parameter plus despiking) denoising (original: all *p* > .63; down-sampled: all *p* > .05).

## Discussion

The present study investigated associations between intrinsic functional brain connectivity dynamics and psychological resilience. More specifically, we explored whether a previously reported negative correlation between resilience and node flexibility as determined using resting state BOLD fMRI (Long et al., 2019) can be replicated in multiband BOLD data with higher temporal resolution (TR = .675 sec). Additionally, we aimed at extending previous findings by including two further metrics of network dynamics, as well as two further questionnaires measuring slightly different theoretical conceptions of resilience. Correlation analyses were performed at different topological scales, i.e., for whole brain, (functional) sub-networks, and nodes. Results did not support the previously reported negative association between node flexibility and resilience, neither in a direct replication matching as closely as possible methodological parameters of the original study, nor when extending the methodology in terms of higher temporal resolution, finer-grained sliding window approach, or further network measures. In these analyses, we found a small number of significant correlations that were distributed rather inconsistently across different measures of brain dynamics and resilience, across atlases, and across sampling schemes: For multiband data (original time resolution; Power264 parcellation), flexibility of a cingulo-opercular node correlated with resilience measured using the CD-RISC questionnaire and promiscuity of a visual node correlated with BRS resilience, however both in the opposite direction relative to Long et al. (2019). For down-sampled data, we observed negative correlations between global degree and BRS and between the degree of a visual node and the RS score (Schaefer100 parcellation). After increasing the spatial resolution to 200 nodes (follow-up analysis), these effects disappeared, and negative resilience correlations emerged for the limbic network (CDRSIC, RS). When using less stringent denoising (as in the original study), we also observed negative correlations with limbic network degree (all three resilience scales), for multiband but not down-sampled data. When applying the AAL atlas, a negative correlation emerged with promiscuity for a subcortical node (pallidum; BRS). Of note, we also observed a borderline significant (*p* = .05) negative correlation between global flexibility and resilience (BRS) in the direct replication. We do not consider this strong support in favor of a replication since we found no similar effects using the Power264 or Schaefer100 atlases or with the other two questionnaires. In sum, these heterogeneous results cannot be considered a successful replication. The fact that the limbic network was involved in these correlations multiple times suggests that its role for resilience should be investigated further. On the other hand, these effects were not consistent, so that they should be treated with caution. Lastly, post-hoc analyses conducted with multiband data and the Schaefer100 parcellation suggest invariance of results against changes in windowing schemes (size, overlap of windows). In the following, we will discuss potential limitations of the present study, which may be important when considering reasons for differences between present and previous results. We then discuss in more depth factors that may account for the failed replication of the results of Long et al. (2019).

### Potential Limitations

Potential limitations of investigating network dynamics in BOLD data with low temporal resolution, as in the original study, have already been discussed in the *Introduction*. When analyzing temporally highly-resolved multiband data, we chose the window size to be equal to 1/f_min_ (Leonardi & Van De Ville, 2015). We cannot rule out that such long window sizes might hamper the sensitivity to small changes, as calculating connectivity matrices over longer periods serves as a smoothing kernel minimizing the ability to resolve TVFC (Vergara et al., 2019). However, this disadvantage of MB-EPI is more than compensated by the ability to acquire more data in the same time, the possibility to investigate faster dynamics, and to thus better disentangle neural and physiological signals (Yang & Lewis, 2021). Furthermore, we showed empirically that node flexibility was largely robust against changes in window size.

Potential limitations may also stem from the questionnaires used: Short versions of two scales (CD-RISC: 10 items, RS: 13 items) were used due to timing constraints, whereas the long (25-items) CD-RISC was used by Long et al. (2019). However, we deem this unproblematic as validity studies show that the short versions can be considered valid and reliable instruments for measuring trait resilience (Chmitorz et al., 2018; Leppert et al., 2008; Sarubin et al., 2015). Our data confirm this, as all questionnaires showed satisfying levels of internal consistency.

### Methodological considerations

Even though the overall direction of correlations between node flexibility and resilience was the same in two previous publications using fMRI (Long et al., 2019) and EEG (Paban et al., 2019), neural localizations partly differed. Adding to this heterogeneity, correlation results in the present study were highly inconsistent (see above) and results thus provide no strong evidence for an association between network dynamics and resilience. This demonstrates the importance of replication attempts also in network neuroscience studies of brain-behavior correlations. To support such work in the future, we have compiled a list of methodological features that we have found important in the present replication study (Table 4).

**Table 4:** Recommendations for individual difference research using dynamic functional connectivity

| **area** | **keyword** | **recommendation** |
| --- | --- | --- |
| methodological | parcellations | test different parcellation schemes and atlases |
|  |  | test different node resolutions to explore stability of effects (currently only for Schaefer atlas possible) |
|  | preprocessing,<br>denoising | optimize preprocessing and denoising<br>strategies to different types of functional<br>connectivity indices |
|  |  | test influence of different denoising<br>pipelines to identify possible relationship<br>between motion and measure of interest |
|  |  | exclude high motion subjects (rather strict<br>than lenient if amount of data allows for) |
|  | sliding-window<br>technique, dynamic<br>functional connectivity | test different windowing schemes (e.g.,<br>size of windows, amount of overlap) |
|  |  | use sufficient amount of data, if possible,<br>e.g., via multiband fMRI (but take into<br>account that acceleration decreases<br>signal-to-noise-ratio) or longer<br>measurements |
|  | multilayer modularity | test different parameter settings |
| psychological | construct of interest | incorporate different measures for (e.g.,<br>two resilience scales) or aim for a<br>complete characterization (i.e., all<br>possible metrics) for the construct of<br>interest, if possible |
|  | brain – construct of<br>interest relationship /<br>confounding variables | motivate in/exclusion of covariates (test<br>both if applicable) |
|  |  | provide descriptive statistics for<br>measures of interest |
|  |  | provide reliability measures (if applicable) |
| replications | method section | provide enough detail to allow for replication attempts |

Note: This brief list of recommendations does not claim to be complete, but rather advocates to always aim at incorporating the most recent findings and empirical evidence from methodological studies.

#### Physiological Noise and Preprocessing

When applying less stringent denoising, a positive correlation between global flexibility and mean framewise displacement (FD) emerged (in multiband time resolution, overlapping windows), suggesting that increased motion inside the scanner may artificially alter functional connectivity and network flexibility. We did not observe any other correlation between dynamic brain measures and motion, suggesting that denoising approaches and their effectiveness may interact with temporal sampling schemes (i.e., MB vs conventional EPI). This underscores the importance of incorporating recent empirical insights regarding denoising strategies for module detection and analyses of network reconfigurations (Lydon-Staley et al., 2018).

#### Sliding Window Technique

Both windowing schemes used in our study (overlapping vs non-overlapping) are among the most-used approaches within the sliding window framework (Iraji et al., 2020). For window-based TVFC analyses, window length is a critical aspect, as it has to balance the ability of robustly estimating TVFC (which benefits from higher numbers of timepoints) against susceptibility to noise (Damaraju et al., 2020; Iraji et al., 2020). Extensive studies have evaluated the impact of different window types, lengths, and overlap on TVFC, however without yet identifying a gold standard (e.g., Abrol et al., 2017; Shakil et al., 2016). At present, lengths between 30s and 100s are commonly used (see Figure S1 in Preti et al., 2017). Within this range, one would however assume that strong effects should be robust against specific analysis choices (compare, e.g., Damaraju et al., 2020 vs. Haimovici et al., 2017). This is supported by our follow-up analyses showing that results did not vary between different window schemes or sizes.

#### Reliability

Recent research suggests that multilayer modularity in particular (Yang et al., 2021) and TVFC in general (Hindriks et al., 2016) are a function of scanning duration and that rs-fMRI sequences around 10 min with standard TRs (as used by Long et al., 2019) are not sufficient for reliable parameter estimation. As reliability strongly influences which correlations are detectable (Hedge et al., 2018), short measurements may be prone to false positive correlations. Yang et al. (2021) propose that at least 20 min of rs-fMRI is needed for reliable multilayer modularity analysis - in that study based on a TR of 1.45 sec and non-overlapping windows of size 100 sec. Here, we measured 10 minutes of rs-fMRI with a TR of about half that length and used a substantially larger number of windows, intentionally selected to allow for reliable multilayer analyses. Yang et al. (2021) also suggest that the standard settings for intra- and interlayer parameters (*γ* = *ω* = 1) in dynamic network measures may not necessarily turn out to be optimal across datasets and across different network measures in terms of reliability. However, when varying the intra- and interlayer parameters beyond the standard setting (in the original/MB data, Schaefer100 atlas; see Supplementary File), we did not observe any significant correlations between node flexibility or promiscuity and resilience scores, except for a single correlation between the flexibility of a visual node and BRS when setting *γ* = 1.4 (*r* = .50, *p* = .02). This correlation, however, points into the opposite direction of what had been reported before (Long et al., 2019). We can thus conclude that the specific choice of intra- and inter-layer parameters has no strong effect on detecting resilience-flexibility correlations.

#### Resilience questionnaires

It is also important that the behavioral variables of interest show satisfying psychometric properties. Internal consistencies for the resilience questionnaires were satisfactory, ranging from .70 to .85. When investigating individual differences, samples should not be too homogeneous on the target measures, as inter-person variability is a precondition for detecting correlations. Even though CD-RISC data appear to be in a somewhat narrower range and more left-skewed than in Long et al. (2019), both the RS-13 and the BRS show sufficient variability (Table 1), which increases confidence that our results are not driven by a lack of heterogeneity. Moreover, resilience scores in our sample were distributed similar to the respective original publications of these measures. It is however difficult to compare our behavioral results directly to the studies by Paban et al. (2019) and Long et al. (2019), as the former did not report descriptive values (but high internal consistency: Cronbachs α = .90), whereas the latter did report descriptive statistics but no psychometrics.

## Conclusion

To summarize, our results do not provide support for the previously reported negative association between node flexibility and psychological resilience. We extended previous studies by including additional measures of functional connectivity dynamics and two further resilience questionnaires and found only weak and inconsistent evidence of associations between network dynamics and resilience, mostly for node degree, a proxy for TVFC, but not for node flexibility or promiscuity. Our study highlights how specific degrees of freedom in the analysis of functional connectivity may influence the presence or absence of effects of interest. This underscores the need for testing the robustness and generalizability of proposed effects via replication.

## Methods

Code and data have been deposited at https://zenodo.org/ under the doi:10.5281/zenodo.5113574 and will be made publicly available after an embargo period that ends December 31, 2024.

### Participants

In total, N = 69 right-handed University students were enrolled in the study, of whom N = 60 completed the study protocol and were included for further analyses. All participants were native speakers of German, right handed, and between 18 and 35 years old; absence of current psychiatric episode was assured with a structured interview (MINI; Sheehan et al., 1998). This sample size was based on considerations of statistical power for investigating across-participant relationships between BOLD activation and cognitive flexibility (see preregistration: https://osf.io/a64jn); the task-free (resting state) fMRI data reported here were acquired together with the pre-registered task-based fMRI experiments. During image quality checks and pre-processing, eight datasets were excluded from further analyses (four due to low quality of questionnaire data and four due to motion artifact; see below for details of exclusion criteria), so that the reported analyses are based on data from 52 participants (26 females, 25 males, 1 diverse; 18-34 years; mean age 24.0 ± 3.7). Using the pwr package in R (Champely et al., 2017), we estimated that our final sample size of N = 52 has a power of > .99 to detect correlational effects of the size reported by Long et al (2019), i.e., of around *r* = .55. All participants provided written informed consent and all procedures were approved by the Ethics Committee of the Department of Psychology of Goethe University Frankfurt, Germany.

### Functional MR Image Acquisition

Resting state functional MRI (rsfMRI) data were collected prior to task-based fMRI (which is not part of the present report; see previous paragraph) on a 3-T Siemens Prisma MR-Scanner equipped with a 32-channel head coil, using a multiband (MB-factor = 4) echo planar imaging (EPI) sequence with the following parameters: 900 volumes (10:14 min), TR = 675 ms, TE = 30 ms, voxel size = 3 mm^3^, flip angle = 60°, FoV = 222 mm, acquisition matrix = 74’74, 40 slices. During the rsfMRI measurement, participants were asked to keep their eyes open and gaze at a white fixation cross, located at the center of a screen (NNL; Nordic Neuro Lab, 40’’, 1920×1080, 60Hz), to stay relaxed and not to think about anything specific. In a separate session, a T1 weighted (T1w) 3D structural MR scan was acquired with a MPRAGE sequence (4:26 min, voxelsize = 1 mm^3^, TR = 1900 ms, TE = 2.52 ms, acquisition matrix = 256×256, 192 slices) for purposes of co-registration between functional and structural data.

### Resilience Questionnaires

The psychological construct ‘resilience’ was quantified using German versions of three self-report questionnaires, the 10 item version of Connor-Davidson Resilience Scale (CD-RISC – 10 items; Connor & Davidson, 2003; German version by Sarubin et al., 2015), the Brief Resilience Scale BRS – 6 items (Smith et al., 2008; German version by Chmitorz et al., 2018), and the Resilience Scale (RS – 13 items; Wagnild & Young, 1993; German version by Leppert et al., 2008). Note that short versions of the questionnaires were selected, as the present study was part of a more extensive study protocol and the short-version have been shown to allow for a time-efficient data collection with comparable validity and reliability (Chmitorz et al., 2018; Leppert et al., 2008; Sarubin et al., 2015). We included the CD-RISC and BRS because of their good ratings in an evaluation study by Windle and colleagues (2011). The RS was additionally chosen to increase comparability with other research, as this questionnaire is frequently used in resilience research. In addition, the selected questionnaires also differ in that CD-RISC and RS define resilience as a personality trait, whereas resilience is understood as an outcome in the BRS (see original publications for more details). Resilience questionnaires were filled out during an informational preparation session for the fMRI measurements and administered online using Unipark software (EFS Survey, Questback GmbH). As completion time of online questionnaires has been identified as the most reliable indicator of data being meaningful or meaningless (Leiner, 2013), we evaluated the quality of questionnaire data using a quality index provided by the Unipark system that compares the completion time of each participant with the average completion time of our sample. As preregistered (https://osf.io/c94y8) participants with an Unipark quality index of .20 or lower were excluded, resulting in exclusion of five participants (see also Leiner, 2013 for a similar criterion). Resilience scores were calculated according to the respective manuals. For each resilience scale, we also assessed internal consistency by calculating Cronbach’s alpha and its 95% confidence interval (CI).

### MR Data Quality Control

Quality of imaging data was assessed using both fMRIPREP’s visual reports as well as MRIQC 0.15.2rc1 (Esteban et al., 2017, 2019). T1w and functional images for each participant were visually checked for signal artifacts, whole brain coverage, and correct alignment between structural and functional data. Following a procedure proposed by Faskowitz and colleagues (2019) functional data were excluded if marked as an outlier (i.e., exceeding 1.5 x the inter-quartile-range either from Q1 or Q3) in more than 50% of the MRIQC quality metrics: *dvars, tsnr, snr, efc, aor, aqi* (see the MRIQC documentation, (Esteban et al., 2017), for more information about these metrics). Given the sensitivity of resting state analyses to movements and given that some of the aforementioned metrics are influenced by motion, we additionally included framewise displacement (FD) as a metric for quantifying motion artifacts (Maknojia et al., 2019). Due to the higher sampling rate of multiband EPI, motion parameters exhibit a high frequency (HF) component resulting from head motion due to respiration (Power et al., 2019) that is usually not observable with standard single-band fMRI (Williams & Van Snellenberg, 2019). With this ‘spurious’ HF motion component, the head appears to be in constant motion and summary measures (such as FD) might be contaminated, leading to exaggerated flagging of ‘bad’ volumes. Note that this applies less to subsequent functional connectivity measures, as those are routinely band-pass filtered, whereas summary measures (e.g., FD) are calculated on raw motion parameters (Gratton et al., 2020; Power et al., 2019; Williams & Van Snellenberg, 2019). We therefore calculated filtered FD (FD_filt_) from Butterworth-filtered raw head movement traces to better isolate ‘true’ head movements (Power et al., 2019). Following Lydon-Staley et al. (2018), we excluded subjects with mean FD_filt_ >0.2 mm or >20 volumes with FD_filt_ >0.25 mm to ensure high data quality. We excluded four subjects based on their FD_filt_, while no subject was excluded based on MRIQC’s metrics.

### Image Preprocessing

T1w and rfMRI images were preprocessed using *fMRIPREP* 20.1.1 (Esteban et al., 2019) which is based on Nipype 1.5.0 (Gorgolewski et al., 2011). A boilerplate text released under a CC0 license describing preprocessing details can be found in the Supplementary File. For further pipeline information, see fMRIPREP’s documentation (Esteban et al., 2019). Due to the use of MB data and the high sampling rate, no slice time correction was applied (see e.g., Glasser et al., 2013). Distortion corrected functional images in T1w space were further denoised using the XPC Engine 1.2.1 (Ciric et al., 2017). We implemented a denoising strategy that has been shown to be relatively effective in mitigating motion artifacts in the study of dynamic functional connectivity, (multilayer) subnetwork detection and measures of module reconfiguration (Lydon-Staley et al., 2018). BOLD data was first demeaned, detrended, and despiked on a voxelwise basis (instead of using more aggressive censoring methods that may result in varying window lengths across participants; (Hutchison et al., 2013) and then temporally filtered with a first-order Butterworth-filter using a passband of 0.01 – 0.08 Hz. These operations were followed by a confound regression, that included (1) six motion estimates derived from fMRIPREPs realignment, (2) mean signals from white matter (WM) and cerebrospinal fluid (CSF), (3) mean global signal, (4) temporal derivatives of these 9 regressors, as well as (5) the quadratic terms of all 18 parameters, resulting in a 36-parameter model to obtain residual BOLD time series. All regressors were also bandpass filtered to avoid reintroducing noise caused by a frequency-dependent mismatch (Hallquist et al., 2013). For the direct replication of Long et al. (2019) and to directly test the influence of different denoising schemes, we also implemented a more lenient 26-paramter denoising approach which included 24 motion parameters following (Friston et al., 1996) together with mean signals from WM and CSF. Within the XCP engine, the Schaefer (resolution: 100, 200; Schaefer et al., 2018), AAL90 (Tzourio-Mazoyer et al., 2002), and Power264 (Power et al., 2011) atlases were transformed to native T1 space and resampled to match the BOLD images (see Supplemental Material for details). For each subject the whole brain was parcellated into distinct regions and a functional time series was extracted for each region, corresponding to the average across all voxels within that region.

### Time Varying Functional Connectivity (TVFC)

All subsequent TVFC analyses were performed with the numpy (1.18.5; Harris et al., 2020) and scipy (1.5.0, SciPy 1.0 Contributors et al., 2020) packages in python 3.8.3.

#### Down-Sampling and window scheme

To directly replicate findings described in Long et. al (2019) and to investigate if different sampling rates (i.e., MB-EPI vs conventional EPI) yield different results, we down-sampled our original resting state BOLD timeseries by a factor of three to resemble a TR of 2 as closely as possible (3 x .675 sec = 2.025 sec). Raw motion traces were also down-sampled accordingly, and FD was calculated without filtering. For the direct replication, we segmented the time series into 15 non-overlapping windows with length of 20 TR (40,5 sec), analogous to the windowing scheme used in Long et al., 2019. As this low number of windows may hamper the ability to investigate brain network dynamics (Hindriks et al., 2016; Lurie et al., 2020; Yang et al., 2021), we additionally used overlapping windows that were shifted by a single timepoint (TR). All windows were tapered with a Hamming filter to reduce potential edge artifacts and to suppress spurious correlations (Shakil et al., 2016; Zalesky et al., 2014), and pairwise Pearson correlation coefficients between all nodes were calculated within each window.

For analyses of the ‘original’ MB data with high temporal resolution, we used a fixed window length of 148 time-points (100 sec), that satisfies the frequency criterion that the length should be at least be equal to 1/f_min_ (Leonardi & Van De Ville, 2015) and allows for a full oscillation of the slowest frequency in the range of 0.01 – 0.08Hz. After down-sampling, sliding window correlations with an adjusted window length of 50 timepoints (101.25 s) and subsequent TVFC and modularity analyses were calculated as described above. To investigate whether the calculated network measures differ between original and down-sampled data and to rule out any idiosyncratic algorithmic behavior (i.e., from multilayer modularity), we compared all three metrics at the nodal level, as these were the starting points for all subsequent measures. For node flexibility and node promiscuity we calculated the spatial similarity (i.e., Spearman correlation coefficient) between mean values per node (average over participants) in the original and down-sampled data, to explore whether nodes behave in comparable manner in both time series. As the down-sampling creates ‘sparser’ data and disrupts the smoothness of the original data (which might result in more abrupt changes in community-assignments), we anticipated a tendency towards higher values for both metrics in the down-sampled data. For node degree, we expected lower values in the down-sampled data, as a recent study by Pedersen et al. (2018) showed that nodal degree decreased with less available data and thus tested this hypothesis using a paired t-test.

We calculated the standard deviation (SD) of each node x node correlation (i.e., node-node connection or edge) over time as a proxy for TVFC (often also referred to as ‘dynamic’ connectivity; Lurie et al., 2020). To test whether these SDs likely reflect ‘true’ dynamics, we benchmarked these estimates against phase randomized surrogate (null) data that preserved auto-correlation, power spectral density and stationary cross correlation of the observed data (Lurie et al., 2020; Prichard & Theiler, 1994; Savva et al., 2019). More precisely, we created 500 surrogates for each subject by phase randomizing the empirical timeseries obtained from the 100 nodes. To preserve the correlative structure between node timeseries, all signals were multiplied by the same uniformly random phase and the SD for each edge was calculated, respectively (see Savva et al., 2019 for a detailed description of phase randomization). The empirically observed SDs were ranked against this null distribution and *p*-values were obtained by dividing the number of times SD_surr_ >= SD_real_ by the number of surrogates. Given that each subject has 4,950 unique connections, FDR-correction (*p* < 0.05) was applied to reduce type-I-errors.

To describe ‘dynamics’ in nodal space, we calculated the nodal degree metric as the binary sum of significant (i.e., ‘dynamic’) connections for each node (Pedersen et al., 2018). Of note, when testing TVFC with surrogate null data, one needs to be cautious as the absence of significance (relative to the null model) does not necessarily indicate the absence of dynamic connectivity. This interpretation depends heavily on the null model applied, and the data may contain meaningful fluctuations relative to other definitions of the null model (see Lurie et al., 2020 for a comprehensive overview).

### Multilayer Modularity

For multilayer modularity, negative correlations in each node x node matrix (i.e., per subject and time window) were set to zero and correlations were z-transformed as done in previous studies (e.g., Finc et al., 2020; Pedersen et al., 2018). We constructed an ordinal multilayer network in which each layer (i.e., time window) represents a weighted adjacency matrix. To assess the spatiotemporal community structure and to track network reconfigurations, each node was linked to itself across layers. To detect communities (i.e., groups of nodes that are more densely connected to one another than to the rest of the network), we used the multilayer counterpart (Mucha et al., 2010) of the modularity function proposed by Newman and Girvan (2004). To optimize the multilayer modularity function, we used an iterative and generative Louvain like algorithm (implemented with code from Jeub et al., https://github.com/GenLouvain/GenLouvain; see Supplementary File for details). The tunable parameters *γ* and *ω* were held constant across layers and set to unity, as had been done in previous studies (e.g., Braun et al., 2015; Yin et al., 2020). As the modularity approach is not deterministic, we repeated it 100 times for each participant and chose the run yielding the highest modularity value, as e.g., done in Finc et al. (2020). We also tested the influence of varying the intra- and interlayer parameters beyond the standard setting in the original data (see Supplementary File) but found no substantial influence on the reported results.

### Time Resolved Analyses of Brain Network Reconfiguration

Within the time-varying community framework, we assessed *node flexibility* as the number of times a node changes its community assignment between adjacent layers, normalized by the total number of possible changes. Node flexibility can be interpreted as a metric that allows to quantify reconfigurations of functional connectivity patterns that a brain region undergoes over time (Braun et al., 2015). To enrich the spatiotemporal description of a node, we further calculated *node promiscuity*, defined as the fraction of communities a node participates in at least once, across all layers. This metric allows to quantify the distribution of a node`s connections over time, e.g., whether high flexibility stems from a switching between two communities or an evenly distributed allegiance to a larger number of different modules (Garcia et al., 2018; Papadopoulos et al., 2016). The higher the nodal promiscuity, the more modules a node participates at least once across time. For each of the three node-specific measures of dynamic network reconfiguration, i.e., flexibility, promiscuity, and degree, we additionally calculated the respective global measure as the average across all nodes, as well as network-specific measures by averaging across all nodes belonging to the respective resting-state sub-networks (RSNs), which varied across atlases.

### Correlation Analyses

All correlational analyses were performed with the pingouin statistics package (0.3.11; Vallat, 2018) in python 3.8.3. To test for associations between resilience and time-varying brain network measures, we calculated partial (Spearman) correlations with age, gender, and FD included as covariates of no interest. Note that unlike Long et al. (2019), years of education was not included as covariate, as all participants were students and years of education were thus not acquired in our study. Correlations were performed at the global, nodal, and network-specific levels for all pairwise combinations between resilience questionnaires (CD-RISC, BRS, RS) and network measures (flexibility, promiscuity, degree). Results were corrected for multiple statistical comparisons using the false discovery rate (*p* < .05; Benjamini & Hochberg, 1995).

### Follow-up Analyses

Lastly, to more directly assess the effect of specific analysis choices on observed correlations between network measures and resilience, we varied a number of such factors systematically, using as ‘standard’ for comparison the analysis pipeline based on MB data parcellated using the Schaefer100 atlas, 36-parameter denoising, and overlapping sliding window scheme. We investigated the effects of including vs. excluding covariates, varying sliding window size (in a frequently used range between 40 and 100 sec in steps of 20 sec; (Preti et al., 2017), and how the two different denoising pipelines affect resilience correlations. Lastly, we also explored within the same parcellation scheme (the Schaefer atlas) the effects of increasing the spatial resolution (i.e., from 100 to 200 nodes).

## Supplementary File 1

- fMRI preprocessing details
- Multilayer Modularity Details
- Correlational results on the nodal level without covariates (Figure S1, S2)
- Node – network assignment in the AAL, Power264 atlas
- Effect of different intra- and interlayer parameters on measures of brain dynamics (Figure S3)
- Effect of different intra- and interlayer parameters on the correlation between measures of brain dynamics and psychological resilience

## Author Contributions

Conceptualization: D.K., C.J.F.

Software: D.K.

Project administration: D.K., C.J.F.

Investigation: D.K.

Data curation: D.K.

Formal analysis: D.K.

Visualization: D.K.

Writing – original draft: D.K., C.J.F.

Writing – review & editing: D.K., C.J.F.

## Acknowledgments

We would like to thank Dr. Cindy Eckart for providing valuable comments on an earlier version of this manuscript and our research assistants for their help in data collection.

## References

Abrol, A., Damaraju, E., Miller, R. L., Stephen, J. M., Claus, E. D., Mayer, A. R., & Calhoun, V. D. (2017). Replicability of time-varying connectivity patterns in large resting state fMRI samples. NeuroImage, 163, 160–176. https://doi.org/10.1016/j.neuroimage.2017.09.020

Bassett, D. S., Wymbs, N. F., Porter, M. A., Mucha, P. J., Carlson, J. M., & Grafton, S. T. (2011). Dynamic reconfiguration of human brain networks during learning. Proceedings of the National Academy of Sciences, 108(18), 7641–7646. https://doi.org/10.1073/pnas.1018985108

Benjamini, Y., & Hochberg, Y. (1995). Controlling the False Discovery Rate: A Practical and Powerful Approach to Multiple Testing. Journal of the Royal Statistical Society. Series B (Methodological), 57(1), 289–300.

Bolsinger, J., Seifritz, E., Kleim, B., & Manoliu, A. (2018). Neuroimaging Correlates of Resilience to Traumatic Events—A Comprehensive Review. Frontiers in Psychiatry, 9, 693. https://doi.org/10.3389/fpsyt.2018.00693

Braun, U., Schäfer, A., Walter, H., Erk, S., Romanczuk-Seiferth, N., Haddad, L., Schweiger, J. I., Grimm, O., Heinz, A., Tost, H., Meyer-Lindenberg, A., & Bassett, D. S. (2015). Dynamic reconfiguration of frontal brain networks during executive cognition in humans. Proceedings of the National Academy of Sciences, 112(37), 11678–11683. https://doi.org/10.1073/pnas.1422487112

Champely, S., Ekstrom, C., Dalgaard, P., Gill, J., Weibelzahl, S., Anandkumar, A., Ford, C., Volvic, R., & De Rosario, H. (2017). pwr: Basic functions for power analysis. https://cran.r-project.org/web/packages/pwr/

Chen, T., Cai, W., Ryali, S., Supekar, K., & Menon, V. (2016). Distinct Global Brain Dynamics and Spatiotemporal Organization of the Salience Network. PLOS Biology, 14(6), e1002469. https://doi.org/10.1371/journal.pbio.1002469

Chmitorz, A., Wenzel, M., Stieglitz, R.-D., Kunzler, A., Bagusat, C., Helmreich, I., Gerlicher, A., Kampa, M., Kubiak, T., Kalisch, R., Lieb, K., & Tüscher, O. (2018a). Population-based validation of a German version of the Brief Resilience Scale. PLOS ONE, 13(2), e0192761. https://doi.org/10.1371/journal.pone.0192761

Chmitorz, A., Wenzel, M., Stieglitz, R.-D., Kunzler, A., Bagusat, C., Helmreich, I., Gerlicher, A., Kampa, M., Kubiak, T., Kalisch, R., Lieb, K., & Tüscher, O. (2018b). Population-based validation of a German version of the Brief Resilience Scale. PLOS ONE, 13(2), e0192761. https://doi.org/10.1371/journal.pone.0192761

Ciric, R., Wolf, D. H., Power, J. D., Roalf, D. R., Baum, G. L., Ruparel, K., Shinohara, R. T., Elliott, M. A., Eickhoff, S. B., Davatzikos, C., Gur, R. C., Gur, R. E., Bassett, D. S., & Satterthwaite, T. D. (2017). Benchmarking of participant-level confound regression strategies for the control of motion artifact in studies of functional connectivity. NeuroImage, 154, 174–187. https://doi.org/10.1016/j.neuroimage.2017.03.020

Connor, K. M., & Davidson, J. R. T. (2003). Development of a new resilience scale: The Connor-Davidson Resilience Scale (CD-RISC). Depression and Anxiety, 18(2), 76–82. https://doi.org/10.1002/da.10113

Dadi, K., Rahim, M., Abraham, A., Chyzhyk, D., Milham, M., Thirion, B., & Varoquaux, G. (2019). Benchmarking functional connectome-based predictive models for resting-state fMRI. NeuroImage, 192, 115–134. https://doi.org/10.1016/j.neuroimage.2019.02.062

Damaraju, E., Tagliazucchi, E., Laufs, H., & Calhoun, V. D. (2020). Connectivity dynamics from wakefulness to sleep. NeuroImage, 220, 117047. https://doi.org/10.1016/j.neuroimage.2020.117047

Dong, D., Wang, Y., Chang, X., Luo, C., & Yao, D. (2018). Dysfunction of Large-Scale Brain Networks in Schizophrenia: A Meta-analysis of Resting-State Functional Connectivity. Schizophrenia Bulletin, 44(1), 168–181. https://doi.org/10.1093/schbul/sbx034

Esteban, O., Birman, D., Schaer, M., Koyejo, O. O., Poldrack, R. A., & Gorgolewski, K. J. (2017). MRIQC: Advancing the automatic prediction of image quality in MRI from unseen sites. PLoS ONE, 12(9). https://doi.org/10.1371/journal.pone.0184661

Esteban, O., Markiewicz, C. J., Blair, R. W., Moodie, C. A., Isik, A. I., Erramuzpe, A., Kent, J. D., Goncalves, M., DuPre, E., Snyder, M., Oya, H., Ghosh, S. S., Wright, J., Durnez, J., Poldrack, R. A., & Gorgolewski, K. J. (2019). fMRIPrep: A robust preprocessing pipeline for functional MRI. Nature Methods, 16(1), 111–116. https://doi.org/10.1038/s41592-018-0235-4

Faskowitz, J., Esfahlani, F. Z., Jo, Y., Sporns, O., & Betzel, R. F. (2019). Edge-centric functional network representations of human cerebral cortex reveal overlapping system-level architecture. BioRxiv, 799924. https://doi.org/10.1101/799924

Finc, K., Bonna, K., He, X., Lydon-Staley, D. M., Kühn, S., Duch, W., & Bassett, D. S. (2020). Dynamic reconfiguration of functional brain networks during working memory training. Nature Communications, 11(1), 2435. https://doi.org/10.1038/s41467-020-15631-z

Friston, K. J., Williams, S., Howard, R., Frackowiak, R. S. J., & Turner, R. (1996). Movement-Related effects in fMRI time-series: Movement Artifacts in fMRI. Magnetic Resonance in Medicine, 35(3). https://doi.org/10.1002/mrm.1910350312

Garcia, J. O., Ashourvan, A., Muldoon, S. F., Vettel, J. M., & Bassett, D. S. (2018). Applications of community detection techniques to brain graphs: Algorithmic considerations and implications for neural function. Proceedings of the IEEE. Institute of Electrical and Electronics Engineers, 106(5), 846–867. https://doi.org/10.1109/JPROC.2017.2786710

Genet, J. J., & Siemer, M. (2011). Flexible control in processing affective and non-affective material predicts individual differences in trait resilience. Cognition & Emotion, 25(2), 380–388. https://doi.org/10.1080/02699931.2010.491647

Gifford, G., Crossley, N., Kempton, M. J., Morgan, S., Dazzan, P., Young, J., & McGuire, P. (2020). Resting state fMRI based multilayer network configuration in patients with schizophrenia. NeuroImage: Clinical, 25, 102169. https://doi.org/10.1016/j.nicl.2020.102169

Glasser, M. F., Sotiropoulos, S. N., Wilson, J. A., Coalson, T. S., Fischl, B., Andersson, J. L., Xu, J., Jbabdi, S., Webster, M., Polimeni, J. R., Van Essen, D. C., & Jenkinson, M. (2013). The Minimal Preprocessing Pipelines for the Human Connectome Project. NeuroImage, 80, 105–124. https://doi.org/10.1016/j.neuroimage.2013.04.127

Gorgolewski, K., Burns, C. D., Madison, C., Clark, D., Halchenko, Y. O., Waskom, M. L., & Ghosh, S. S. (2011). Nipype: A Flexible, Lightweight and Extensible Neuroimaging Data Processing Framework in Python. Frontiers in Neuroinformatics, 5. https://doi.org/10.3389/fninf.2011.00013

Gratton, C., Dworetsky, A., Coalson, R. S., Adeyemo, B., Laumann, T. O., Wig, G. S., Kong, T. S., Gratton, G., Fabiani, M., Barch, D. M., Tranel, D., Miranda-Dominguez, O., Fair, D. A., Dosenbach, N. U. F., Snyder, A. Z., Perlmutter, J. S., Petersen, S. E., & Campbell, M. C. (2020). Removal of high frequency contamination from motion estimates in single-band fMRI saves data without biasing functional connectivity. bioRxiv, 837161. https://doi.org/10.1101/837161

Haimovici, A., Tagliazucchi, E., Balenzuela, P., & Laufs, H. (2017). On wakefulness fluctuations as a source of BOLD functional connectivity dynamics. Scientific Reports, 7(1), 5908. https://doi.org/10.1038/s41598-017-06389-4

Hallquist, M. N., Hwang, K., & Luna, B. (2013). The nuisance of nuisance regression: Spectral misspecification in a common approach to resting-state fMRI preprocessing reintroduces noise and obscures functional connectivity. NeuroImage, 82, 208–225. https://doi.org/10.1016/j.neuroimage.2013.05.116

Harlalka, V., Bapi, R. S., Vinod, P. K., & Roy, D. (2019). Atypical Flexibility in Dynamic Functional Connectivity Quantifies the Severity in Autism Spectrum Disorder. Frontiers in Human Neuroscience, 13, 6. https://doi.org/10.3389/fnhum.2019.00006

Harris, C. R., Millman, K. J., van der Walt, S. J., Gommers, R., Virtanen, P., Cournapeau, D., Wieser, E., Taylor, J., Berg, S., Smith, N. J., Kern, R., Picus, M., Hoyer, S., van Kerkwijk, M. H., Brett, M., Haldane, A., del Río, J. F., Wiebe, M., Peterson, P., … Oliphant, T. E. (2020). Array programming with NumPy. Nature, 585(7825), 357–362. https://doi.org/10.1038/s41586-020-2649-2

Hedge, C., Powell, G., & Sumner, P. (2018). The reliability paradox: Why robust cognitive tasks do not produce reliable individual differences. Behavior Research Methods, 50(3), 1166–1186. https://doi.org/10.3758/s13428-017-0935-1

Hindriks, R., Adhikari, M. H., Murayama, Y., Ganzetti, M., Mantini, D., Logothetis, N. K., & Deco, G. (2016). Can sliding-window correlations reveal dynamic functional connectivity in resting-state fMRIã NeuroImage, 127, 242–256. https://doi.org/10.1016/j.neuroimage.2015.11.055

Hutchison, R. M., Womelsdorf, T., Allen, E. A., Bandettini, P. A., Calhoun, V. D., Corbetta, M., Della Penna, S., Duyn, J. H., Glover, G. H., Gonzalez-Castillo, J., Handwerker, D. A., Keilholz, S., Kiviniemi, V., Leopold, D. A., de Pasquale, F., Sporns, O., Walter, M., & Chang, C. (2013). Dynamic functional connectivity: Promise, issues, and interpretations. NeuroImage, 80, 360–378. https://doi.org/10.1016/j.neuroimage.2013.05.079

Iraji, A., Faghiri, A., Lewis, N., Fu, Z., Rachakonda, S., & Calhoun, V. D. (2020). Tools of the trade: Estimating time-varying connectivity patterns from fMRI data. Social Cognitive and Affective Neuroscience, saa114. https://doi.org/10.1093/scan/nsaa114

Jin, C., Jia, H., Lanka, P., Rangaprakash, D., Li, L., Liu, T., Hu, X., & Deshpande, G. (2017). Dynamic brain connectivity is a better predictor of PTSD than static connectivity. Human Brain Mapping, 38(9), 4479–4496. https://doi.org/10.1002/hbm.23676

Kashdan, T. B., & Rottenberg, J. (2010). Psychological flexibility as a fundamental aspect of health. Clinical Psychology Review, 30(7), 865–878. https://doi.org/10.1016/j.cpr.2010.03.001

Leiner, D. J. (2013). Too Fast, Too Straight, Too Weird: Post Hoc Identification of Meaningless Data in Internet Surveys. SSRN Electronic Journal. https://doi.org/10.2139/ssrn.2361661

Leonardi, N., & Van De Ville, D. (2015a). On spurious and real fluctuations of dynamic functional connectivity during rest. NeuroImage, 104, 430–436. https://doi.org/10.1016/j.neuroimage.2014.09.007

Leonardi, N., & Van De Ville, D. (2015b). On spurious and real fluctuations of dynamic functional connectivity during rest. NeuroImage, 104, 430–436. https://doi.org/10.1016/j.neuroimage.2014.09.007

Leppert, K., Koch, B., Brähler, E., & Strauß, B. (2008). Die Resilienzskala (RS) – Überprüfung der Langform RS-25 und einer Kurzform RS-13.

Li, S., Hu, N., Zhang, W., Tao, B., Dai, J., Gong, Y., Tan, Y., Cai, D., & Lui, S. (2019). Dysconnectivity of Multiple Brain Networks in Schizophrenia: A Meta-Analysis of Resting-State Functional Connectivity. Frontiers in Psychiatry, 10, 482. https://doi.org/10.3389/fpsyt.2019.00482

Long, Y., Chen, C., Deng, M., Huang, X., Tan, W., Zhang, L., Fan, Z., & Liu, Z. (2019). Psychological resilience negatively correlates with resting-state brain network flexibility in young healthy adults: A dynamic functional magnetic resonance imaging study. Annals of Translational Medicine, 7(24), 809. https://doi.org/10.21037/atm.2019.12.45

Lurie, D. J., Kessler, D., Bassett, D. S., Betzel, R. F., Breakspear, M., Kheilholz, S., Kucyi, A., Liégeois, R., Lindquist, M. A., McIntosh, A. R., Poldrack, R. A., Shine, J. M., Thompson, W. H., Bielczyk, N. Z., Douw, L., Kraft, D., Miller, R. L., Muthuraman, M., Pasquini, L., … Calhoun, V. D. (2020). Questions and controversies in the study of time-varying functional connectivity in resting fMRI. Network Neuroscience, 4(1), 30–69. https://doi.org/10.1162/netn_a_00116

Lydon-Staley, D. M., Ciric, R., Satterthwaite, T. D., & Bassett, D. S. (2018). Evaluation of confound regression strategies for the mitigation of micromovement artifact in studies of dynamic resting-state functional connectivity and multilayer network modularity. Network Neuroscience, 1–28. https://doi.org/10.1162/netn_a_00071

Maknojia, S., Churchill, N. W., Schweizer, T. A., & Graham, S. J. (2019). Resting State fMRI: Going Through the Motions. Frontiers in Neuroscience, 13, 825. https://doi.org/10.3389/fnins.2019.00825

Menon, V. (2011). Large-scale brain networks and psychopathology: A unifying triple network model. Trends in Cognitive Sciences, 15(10), 483–506. https://doi.org/10.1016/j.tics.2011.08.003

Mucha, P. J., Richardson, T., Macon, K., Porter, M. A., & Onnela, J.-P. (2010). Community Structure in Time-Dependent, Multiscale, and Multiplex Networks. Science, 328(5980), 876–878. https://doi.org/10.1126/science.1184819

Mulders, P. C., van Eijndhoven, P. F., Schene, A. H., Beckmann, C. F., & Tendolkar, I. (2015). Resting-state functional connectivity in major depressive disorder: A review. Neuroscience & Biobehavioral Reviews, 56, 330–344. https://doi.org/10.1016/j.neubiorev.2015.07.014

Muldoon, S. F., & Bassett, D. S. (2016). Network and Multilayer Network Approaches to Understanding Human Brain Dynamics. Philosophy of Science, 83(5), 710–720. https://doi.org/10.1086/687857

Newman, M. E. J., & Girvan, M. (2004). Finding and evaluating community structure in networks. Physical Review E, 69(2), 026113. https://doi.org/10.1103/PhysRevE.69.026113

Nolen-Hoeksema, S., Wisco, B. E., & Lyubomirsky, S. (2008). Rethinking Rumination. Perspectives on Psychological Science: A Journal of the Association for Psychological Science, 3(5), 400–424. https://doi.org/10.1111/j.1745-6924.2008.00088.x

Paban, V., Modolo, J., Mheich, A., & Hassan, M. (2019). Psychological resilience correlates with EEG source-space brain network flexibility. Network Neuroscience, 3(2), 539–550. https://doi.org/10.1162/netn_a_00079

Papadopoulos, L., Puckett, J. G., Daniels, K. E., & Bassett, D. S. (2016). Evolution of network architecture in a granular material under compression. Physical Review. E, 94(3–1), 032908. https://doi.org/10.1103/PhysRevE.94.032908

Parsons, S., Kruijt, A.-W., & Fox, E. (2016). A Cognitive Model of Psychological Resilience. Journal of Experimental Psychopathology, 7(3), 296–310. https://doi.org/10.5127/jep.053415

Pedersen, M., Zalesky, A., Omidvarnia, A., & Jackson, G. D. (2018). Multilayer network switching rate predicts brain performance. Proceedings of the National Academy of Sciences, 115(52), 13376–13381. https://doi.org/10.1073/pnas.1814785115

Power, J. D., Cohen, A. L., Nelson, S. M., Wig, G. S., Barnes, K. A., Church, J. A., Vogel, A. C., Laumann, T. O., Miezin, F. M., Schlaggar, B. L., & Petersen, S. E. (2011). Functional network organization of the human brain. Neuron, 72(4), 665–678. https://doi.org/10.1016/j.neuron.2011.09.006

Power, J. D., Lynch, C. J., Silver, B. M., Dubin, M. J., Martin, A., & Jones, R. M. (2019). Distinctions among real and apparent respiratory motions in human fMRI data. NeuroImage, 201, 116041. https://doi.org/10.1016/j.neuroimage.2019.116041

Preti, M. G., Bolton, T. A., & Van De Ville, D. (2017). The dynamic functional connectome: State-of-the-art and perspectives. NeuroImage, 160, 41–54. https://doi.org/10.1016/j.neuroimage.2016.12.061

Prichard, D., & Theiler, J. (1994). Generating surrogate data for time series with several simultaneously measured variables. Physical Review Letters, 73(7), 951–954. https://doi.org/10.1103/PhysRevLett.73.951

Sarubin, N., Gutt, D., Giegling, I., Bühner, M., Hilbert, S., Krähenmann, O., Wolf, M., Jobst, A., Sabaß, L., Rujescu, D., Falkai, P., & Padberg, F. (2015). Erste Analyse der psychometrischen Eigenschaften und Struktur der deutschsprachigen 10-und 25-Item Version der Connor-Davidson Resilience Scale (CD-RISC). Zeitschrift für Gesundheitspsychologie, 23(3), 112–122. https://doi.org/10.1026/0943-8149/a000142

Savva, A. D., Mitsis, G. D., & Matsopoulos, G. K. (2019). Assessment of dynamic functional connectivity in resting-state fMRI using the sliding window technique. Brain and Behavior, 9(4), e01255. https://doi.org/10.1002/brb3.1255

Schaefer, A., Kong, R., Gordon, E. M., Laumann, T. O., Zuo, X.-N., Holmes, A. J., Eickhoff, S. B., & Yeo, B. T. T. (2018). Local-Global Parcellation of the Human Cerebral Cortex from Intrinsic Functional Connectivity MRI. Cerebral Cortex (New York, N.Y.: 1991), 28(9), 3095–3114. https://doi.org/10.1093/cercor/bhx179

SciPy 1.0 Contributors, Virtanen, P., Gommers, R., Oliphant, T. E., Haberland, M., Reddy, T., Cournapeau, D., Burovski, E., Peterson, P., Weckesser, W., Bright, J., van der Walt, S. J., Brett, M., Wilson, J., Millman, K. J., Mayorov, N., Nelson, A. R. J., Jones, E., Kern, R., … van Mulbregt, P. (2020). SciPy 1.0: Fundamental algorithms for scientific computing in Python. Nature Methods, 17(3), 261–272. https://doi.org/10.1038/s41592-019-0686-2

Shakil, S., Lee, C.-H., & Keilholz, S. D. (2016). Evaluation of sliding window correlation performance for characterizing dynamic functional connectivity and brain states. NeuroImage, 133, 111–128. https://doi.org/10.1016/j.neuroimage.2016.02.074

Sheehan, D. V., Lecrubier, Y., Sheehan, K. H., Amorim, P., Janavs, J., Weiller, E., Hergueta, T., Baker, R., & Dunbar, G. C. (1998). The Mini-International Neuropsychiatric Interview (M.I.N.I.): The development and validation of a structured diagnostic psychiatric interview for DSM-IV and ICD-10. The Journal of Clinical Psychiatry, 59 Suppl 20, 22–33;quiz 34-57.

Smith, B. W., Dalen, J., Wiggins, K., Tooley, E., Christopher, P., & Bernard, J. (2008). The brief resilience scale: Assessing the ability to bounce back. International Journal of Behavioral Medicine, 15(3), 194–200. https://doi.org/10.1080/10705500802222972

Southwick, S. M., & Charney, D. S. (2012). The Science of Resilience: Implications for the Prevention and Treatment of Depression. Science, 338(6103), 79–82. https://doi.org/10.1126/science.1222942

Tzourio-Mazoyer, N., Landeau, B., Papathanassiou, D., Crivello, F., Etard, O., Delcroix, N., Mazoyer, B., & Joliot, M. (2002). Automated anatomical labeling of activations in SPM using a macroscopic anatomical parcellation of the MNI MRI single-subject brain. NeuroImage, 15(1), 273–289. https://doi.org/10.1006/nimg.2001.0978

Vallat, R. (2018). Pingouin: Statistics in Python. Journal of Open Source Software, 3(31), 1026. https://doi.org/10.21105/joss.01026

Vergara, V. M., Abrol, A., & Calhoun, V. D. (2019). An average sliding window correlation method for dynamic functional connectivity. Human Brain Mapping, 40(7), 2089–2103. https://doi.org/10.1002/hbm.24509

Wagnild, G. M., & Young, H. M. (1993). Development and psychometric evaluation of the Resilience Scale. Journal of Nursing Measurement, 1(2), 165–178.

Williams, J. C., & Van Snellenberg, J. X. (2019). Motion denoising of multiband resting state functional connectivity MRI data: An improved volume censoring method [Preprint]. Neuroscience. https://doi.org/10.1101/860635

Windle, G., Bennett, K. M., & Noyes, J. (2011). A methodological review of resilience measurement scales. Health and Quality of Life Outcomes, 9, 8. https://doi.org/10.1186/1477-7525-9-8

Yang, Z., & Lewis, L. D. (2021). Imaging the temporal dynamics of brain states with highly sampled fMRI. Current Opinion in Behavioral Sciences, 40, 87–95. https://doi.org/10.1016/j.cobeha.2021.02.005

Yang, Z., Telesford, Q. K., Franco, A. R., Lim, R., Gu, S., Xu, T., Ai, L., Castellanos, F. X., Yan, C.-G., Colcombe, S., & Milham, M. P. (2021). Measurement reliability for individual differences in multilayer network dynamics: Cautions and considerations. NeuroImage, 225, 117489. https://doi.org/10.1016/j.neuroimage.2020.117489

Yin, W., Li, T., Hung, S.-C., Zhang, H., Wang, L., Shen, D., Zhu, H., Mucha, P. J., Cohen, J. R., & Lin, W. (2020). The emergence of a functionally flexible brain during early infancy. Proceedings of the National Academy of Sciences, 202002645. https://doi.org/10.1073/pnas.2002645117

Zalesky, A., Fornito, A., Cocchi, L., Gollo, L. L., & Breakspear, M. (2014). Time-resolved resting-state brain networks. Proceedings of the National Academy of Sciences, 111(28), 10341–10346. https://doi.org/10.1073/pnas.1400181111

