## Supplemental File 1 for "Probing the association between resting state brain network dynamics and psychological resilience"

Department of Psychology, Goethe University Frankfurt, Theodor-W.-Adorno-Platz 6, Frankfurt am Main 60323, Germany

**Supplementary File 1**:

- fMRI preprocessing details
- Multilayer Modularity Details
- Correlational results on the nodal level without covariates (Figure S1, S2)
- Node – network assignment in the AAL, Power264 atlas
- Effect of different intra- and interlayer parameters on measures of brain dynamics (Figure S3)
- Effect of different intra- and interlayer parameters on the correlation between measures of brain dynamics and psychological resilience

**Supplementary File 1**

### MRI data preprocessing

After reconstruction BOLD fMRI data were organized using the Brain Imaging Data Structure (BIDS; Gorgolewski et al., 2016) and preprocessed using *fMRIPrep* version 20.1.1 (Esteban et al., 2018a, 2018b; RRID:SCR_016216) which is based on *Nipype* 1.5.0 (Gorgolewski et al., 2011; Gorgolewski et al., 2018; RRID:SCR_002502). Many internal operations of *fMRIPrep* use *Nilearn* 0.6.2 (Abraham et al., 2014; RRID:SCR_001362) mostly within the functional processing workflow; all other software tools used are specified below. For more details of the pipeline see the section corresponding to workflows in *fMRIPrep*'s documentation. In the following, preprocessing is described in more detail by text that was automatically generated by the fMRIPrep software and then manually edited with respect to readability.

#### *Anatomical data preprocessing*

One T1-weighted (T1w) image was found within the input BIDS dataset. The T1-weighted (T1w) image was corrected for intensity non-uniformity (INU) with N4BiasFieldCorrection (Tustison et al., 2010) distributed with ANTs 2.2.0 (Avants et al., 2008; RRID:SCR_004757) and used as T1w-reference throughout the workflow. The T1w-reference was then skull-stripped with a *Nipype* implementation of the antsBrainExtraction.sh workflow (from ANTs 2.2.0) using OASIS30ANTs as target template. Brain tissue segmentation of cerebrospinal fluid (CSF), white-matter (WM), and gray-matter (GM) was performed on the brain-extracted T1w using FAST (FSL 5.0.9; RRID:SCR_002823; Zhang, Brady. & Smith, 2001). Brain surfaces were reconstructed using recon-all (FreeSurfer 6.0.1; RRID:SCR_001847; Dale, Fischl, & Sereno, 1999) and the brain mask estimated previously was refined with a custom variation of the method to reconcile ANTs-derived and FreeSurfer-derived segmentations of the cortical gray-matter of Mindboggle (RRID:SCR_002438; Klein et al., 2017). Volume-based spatial normalization to the *ICBM 152 Nonlinear Asymmetrical template version 2009c* (Fonov et al., 2009, RRID:SCR_008796; TemplateFlow ID: MNI152NLin2009cAsym) standard space was performed through nonlinear registration with antsRegistration (ANTs 2.2.0) using brain-extracted versions of both the T1w reference and the T1w template.

#### *Functional data preprocessing*

For each subject's BOLD run the following preprocessing was performed: First, a reference volume and its skull-stripped version were generated using a custom methodology of *fMRIPrep*: Head-motion parameters with respect to the BOLD reference (transformation matrices and six corresponding rotation and translation parameters) were estimated before any spatiotemporal filtering using MCFLIRT (FSL 5.0.9; Jenkinson et al., 2002). A B0-nonuniformity map (or *fieldmap*) was estimated based on a phase-difference map calculated with a dual-echo GRE (gradient-recall echo) sequence processed with a custom workflow of *Susceptibility Distortion Correction workFlows* (*SDCFlows;* Markiewicz et al., 2020) inspired by the epidewarp.fsl script and further improvements in Human Connectome Project Pipelines (Glasser et al., 2013). The *fieldmap* was then co-registered to the target EPI (echo-planar imaging) reference run and converted to a displacements field map (amenable to registration tools such as ANTs) with FSL's FUGUE and other *SDCflows* tools. Based on the estimated susceptibility distortion a corrected EPI (echo-planar imaging) reference was calculated for a more accurate co-registration with the anatomical reference. The BOLD reference was then co-registered to the T1w reference using bbregister (FreeSurfer) which implements boundary-based registration (Greve & Fischl, 2009). Co-registration was configured with six degrees of freedom. The BOLD time-series were resampled onto FreeSurfer's the *fsnative* and *fsaverage* surfaces. The BOLD time-series were resampled onto their original native space by applying a single composite transform to correct for head-motion and susceptibility distortions. These resampled BOLD time-series will be referred to as *preprocessed BOLD in original space* or just *preprocessed BOLD*.

The BOLD time-series were resampled into standard space generating a *preprocessed BOLD run in MNI152NLin2009cAsym space*. First a reference volume and its skull-stripped version were generated using a custom methodology of *fMRIPrep*: Several confounding time-series were calculated based on the *preprocessed BOLD*, i.e., framewise displacement (FD), derivative of RMS variance over voxels (DVARS), and three region-wise global signals. FD was computed using two formulations following Power (absolute sum of relative motions; Power et al., 2014) and Jenkinson (relative root mean square displacement between affines; Jenkinson et al., 2002). FD and DVARS were calculated for the functional run using their implementations in *Nipype* (following the definitions by Power et al., 2014). The three global signals were extracted within the CSF, WM, and whole-brain masks. Additionally, a set of physiological regressors were extracted to allow for component-based noise correction (*CompCor*; Behzadi et al., 2007). Principal components are estimated after high-pass filtering the *preprocessed BOLD* time-series (using a discrete cosine filter with 128s cut-off) for the two *CompCor* variants: temporal (tCompCor) and anatomical (aCompCor). TCompCor components are then calculated from the top 5% variable voxels within a mask covering the subcortical regions. This subcortical mask is obtained by heavily eroding the brain mask, which ensures it does not include cortical GM regions. For aCompCor, components are calculated within the intersection of the aforementioned mask and the union of CSF and WM masks calculated in T1w space after their projection to the native space of the functional run (using the inverse BOLD-to-T1w transformation). Components are also calculated separately within the WM and CSF masks. For each CompCor decomposition, the *k* components with the largest singular values are retained such that the retained components' time series are sufficient to explain 50 percent of variance across the nuisance mask (CSF, WM, combined, or temporal). The remaining components are dropped from consideration. The head-motion estimates calculated in the correction step were also placed within the corresponding confounds file. The confound time series derived from head motion estimates and global signals were expanded with the inclusion of temporal derivatives and quadratic terms for each (Satterthwaite et al., 2013). All resamplings can be performed with *a single interpolation step* by composing all the pertinent transformations (i.e., head-motion transform matrices. susceptibility distortion correction, and co-registrations to anatomical and output spaces). Gridded (volumetric) resamplings were performed using antsApplyTransforms (ANTs) configured with Lanczos interpolation to minimize the smoothing effects of other kernels (Lanczos, 1964). Non-gridded (surface) resamplings were performed using mri_vol2surf (FreeSurfer).

**Multilayer Modularity details**

In general, community detection and node assignment can be solved algorithmically by maximizing the modularity quality function Q. Q allows to quantify the goodness of fit of a certain partition, with high values indicating high quality partitions (Furtunato, 2010). Following the idea that random graphs do not show a cluster structure, the quality function compares the number of intracommunity edges to what would be expected at random (i.e., under a null model), to quantify communities (Fortunato, 2010; Mucha et al., 2010). The quality function Q can be formalized as follows:

$$Q = \sum_{ij} \left(A_{ij} - P_{ij}\right)\delta\left(M_i, M_j\right) \quad (1)$$

where $A_{ij}$ and $P_{ij}$ are the observed and expected weights of the connection between node *i* and node *j*. Here, $A_{ij}$ is an element of the adjacency matrix $\mathbf{A}$ and $P_{ij}$ refers to a specific null model. The Kronecker delta function $\delta\left(M_i, M_j\right)$ equals 1 if node *i* and *j* are assigned to the same module (M) and 0 if otherwise. Noteworthy, the modularity function suffers from a "resolution limit", meaning that communities that are rather small compared to the whole graph may be undetectable (Fortunato & Barthelemy, 2007). To overcome this caveat, one may introduce a structural resolution parameter $\gamma$ that weights the null model. Optimizing $Q(\gamma)$ with large values of $\gamma$ leads to the detection of many small communities, whereas small values of $\gamma$ result in few large communities. Here, we used an expanded modularity function that can be used with multilayer networks, formalized as follows:

$$Q(\gamma, \omega) = \frac{1}{2\mu} \sum_{ijsr} \left[\left(A_{ijs} - \gamma_s \frac{k_{is}k_{js}}{2m_s}\right)\delta\left(M_{is}, M_{js}\right) + \delta_{ij}\, \omega_{jsr}\right] \delta\left(M_{is}, M_{jr}\right) \quad (2)$$

where $A_{ijs}$ describes elements of $\mathbf{A}$ of layer (time window) s and $\frac{k_{is}k_{js}}{2m_s}$ denotes the Newman-Girvan configuration null model (i.e., the expected connectivity strength at chance level, with k = node degree at layer s; m = sum of all edges at layer s) that is scaled by the resolution parameter $\gamma_s$ of layer s, $\delta\left(M_{is}, M_{js}\right)$ and $\delta\left(M_{is}, M_{jr}\right)$ = 1 if node *i* and node *j* are in the same module (M) or 0 if they do not belong to the same module (here $M_{is}$ depicts the community assignment of node *i* in layer *s*; $\omega_{jsr}$ is the interlayer coupling parameter connecting node *j* in layer s to itself in layer r, defining the similarity of communities across layers with large values of $\omega_{jsr}$ leading to increased homogeneity across layers, as nodes will prefer to stay in one community (Vaiana & Muldoon, 2018).

### Correlation results on a nodal level without covariates.

Figures S1 and S2 depict the correlations between brain network measures on a nodal level and resilience questionnaires calculated without covariates. These figures can be compared to Figure 4 and Figure 5 of the original manuscript, which show the same calculations with inclusion of covariates. Results are described in more detail in the original manuscript.

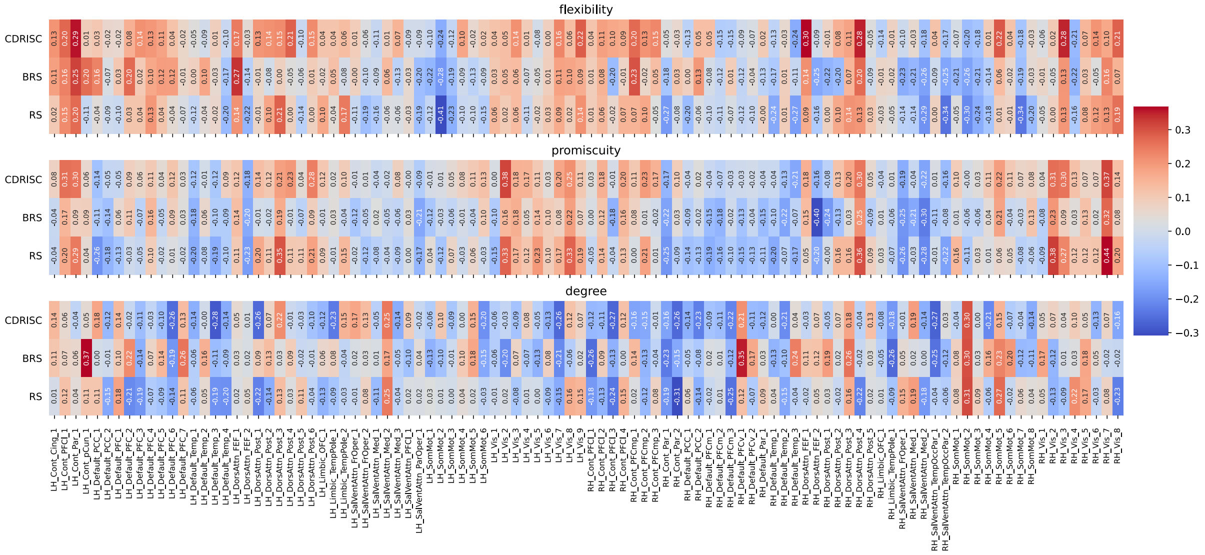


**Figure S1.** Correlations between network measures and resilience at the nodal level for the original (multiband) data (TR = .675 sec), calculated without covariates. Node names are depicted on the x-axis ticks; for a detailed list see https://bit.ly/3yvOBwz. LH = left hemisphere, RH = right hemisphere.

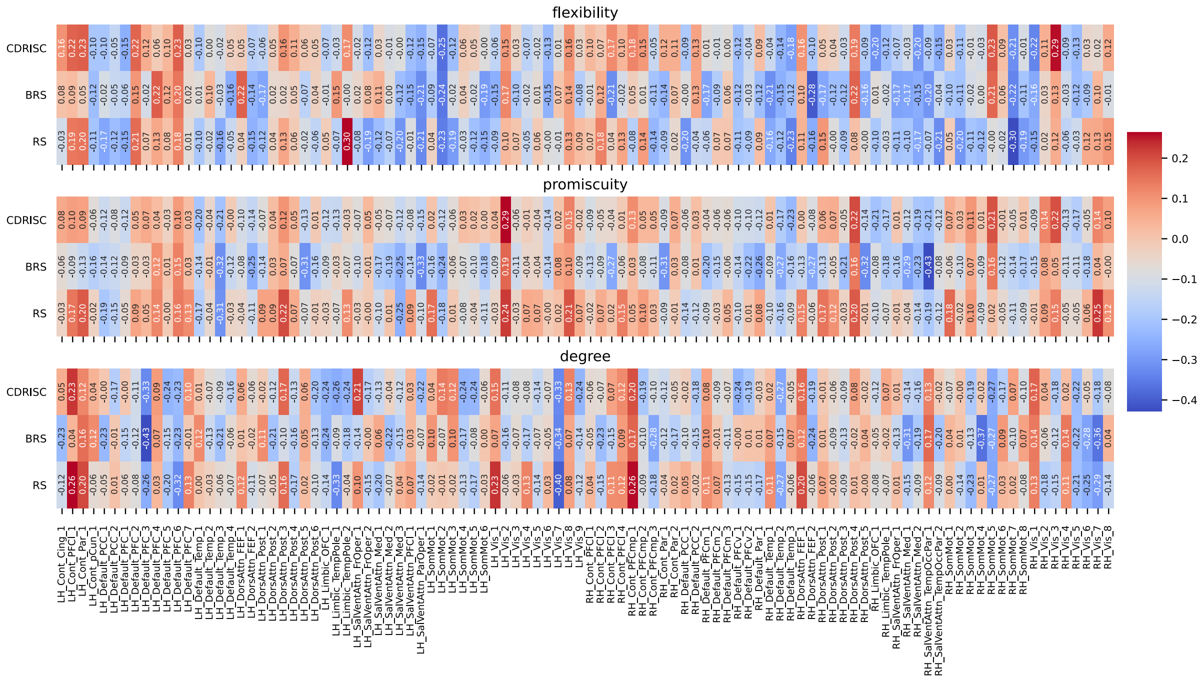


**Figure S2**. Correlations between network measures and resilience at the nodal level for the down-sampled data (TR = 2.025 sec) without the inclusion of covariates. Node names are depicted on the x-axis ticks; for a detailed list see https://bit.ly/3yvOBwz. LH = left hemisphere, RH = right hemisphere.

### Node – network assignment in the AAL, Power264 atlas

To additionally test whether results reported here might have been driven by the specific parcellation we used (i.e., Schaefer 100), we repeated the pre-processing and denoising pipeline with two additional atlases, i.e., the AAL parcellation with 90 regions (Tzourio-Mazoyer et al., 2002) and the atlas of (Power et al., 2011) with 264 regions, both of which had been used in the study by Long and colleagues (2019). For the AAL90 atlas, we referred to the node – RSN assignment provided by Long et al. (2019) in their Table S1. However, according to that Table, some nodes were assigned to multiple subnetworks (e.g., node middle frontal gyrus), which we avoided. Accordingly, our subnetwork assignment may vary in details from the one used in Long et al. (2019). Taking Long et al.'s (2019) Table S1 as a starting point, we assigned each node to the subnetwork which appears first in the respective column, leading to a total number of eight unique networks, i.e., 'Sensorimotor', 'Visual', 'Salience', 'Auditory', 'Subcortical', 'Frontoparietal', 'Default-mode', and 'Cingulo-opercular'. 12 nodes were not assigned to any sub-network (i.e.,'None'), including the amygdala and hippocampus. For the Power atlas (nodes 1, 3, 8, 9, 119, 126, 183, and 184) from further analyses, as some participants were lacking BOLD signal for these nodes. The remaining nodes were assigned to one of the following resting state networks (RSN): 'somatomotor hand', 'somatomotor mouth', 'cingulo-opercular', 'auditory', 'default mode', 'memory retrieval', 'visual', frontoparietal', 'salience', 'subcortical', 'ventral attention', 'dorsal attention', or 'cerebellar'. Analogous to the AAL atlas, 21 nodes that could not be assigned to any of these categories were sorted into a 'None' category. For both parcellations, we again calculated partial correlations between dynamic brain measures and the respective resilience questionnaires on a nodal, RSN, and global scale.

**Effect of different intra- and interlayer parameters on measures of brain dynamics**

The community detection in multilayer networks is tuned by the intra- and inter-layer parameters $\gamma$ and $\omega$, respectively. Whereas the intralayer parameter $\gamma$ determines how many communities will be detected (i.e., larger values leading to the detection of smaller communities) at a given timepoint, the interlayer resolution parameter $\omega$ scales the network dynamics over time, i.e., larger values leading to increased homogeneity over time and nodes consequently tend to switch communities less often (Vaiana & Muldoon, 2018). We here investigate the stability of our correlation results for neural flexibility and promiscuity measures, by systematically varying these parameters. In line with previous studies (e.g., Pedersen et al., 2018; Yin et al., 2020) we considered intralayer parameters ($\gamma$) from 0.8 to 1.4 and interlayer parameters ($\omega$) from 0.4 to 1.0 in steps of 0.2, while fixing the respective other parameter to 1 (using MB data with TR = .675 and the Schaefer100 atlas). The detection of smaller communities (i.e., larger values of $\gamma$) led to higher nodal flexibility (see Figure S3), whereas it led to smaller nodal promiscuity (i.e., nodes tend to change their community assignments more often over time, but these interactions are not distributed across all other modules but appear to be rather limited to a small number of communities. The spatial similarity (i.e., correlations) of nodal flexibility values (range $r$ = .52 to .92, all $p$ < .0001) and promiscuity (range $r$ = .40 to .92, all $p$ < .0001) across the range of parameters studied suggest a moderate to good correspondence across values (see Figure XX). Across layers, lower values of the interlayer parameter $\omega$ led to increased node flexibility, whereas node promiscuity showed the opposite pattern, i.e., decreasing values with lower omega (see Figure S3). However, despite numerical changes, the spatial similarity of node flexibility ($r$ > .95, $p$ < .0001) and promiscuity ($r$ > .96, $p$ < .0001) for all parameters of $\omega$ was high. Both patterns suggest that nodes switch between communities more often over time, but that this increase is driven by a constant switching between a small number of communities and not a broad interaction with various communities (i.e., simultaneous low promiscuity with high flexibility). More generally, these results demonstrate relative robustness of node flexibility and node promiscuity measures against variations in the parameters gamma and omega.

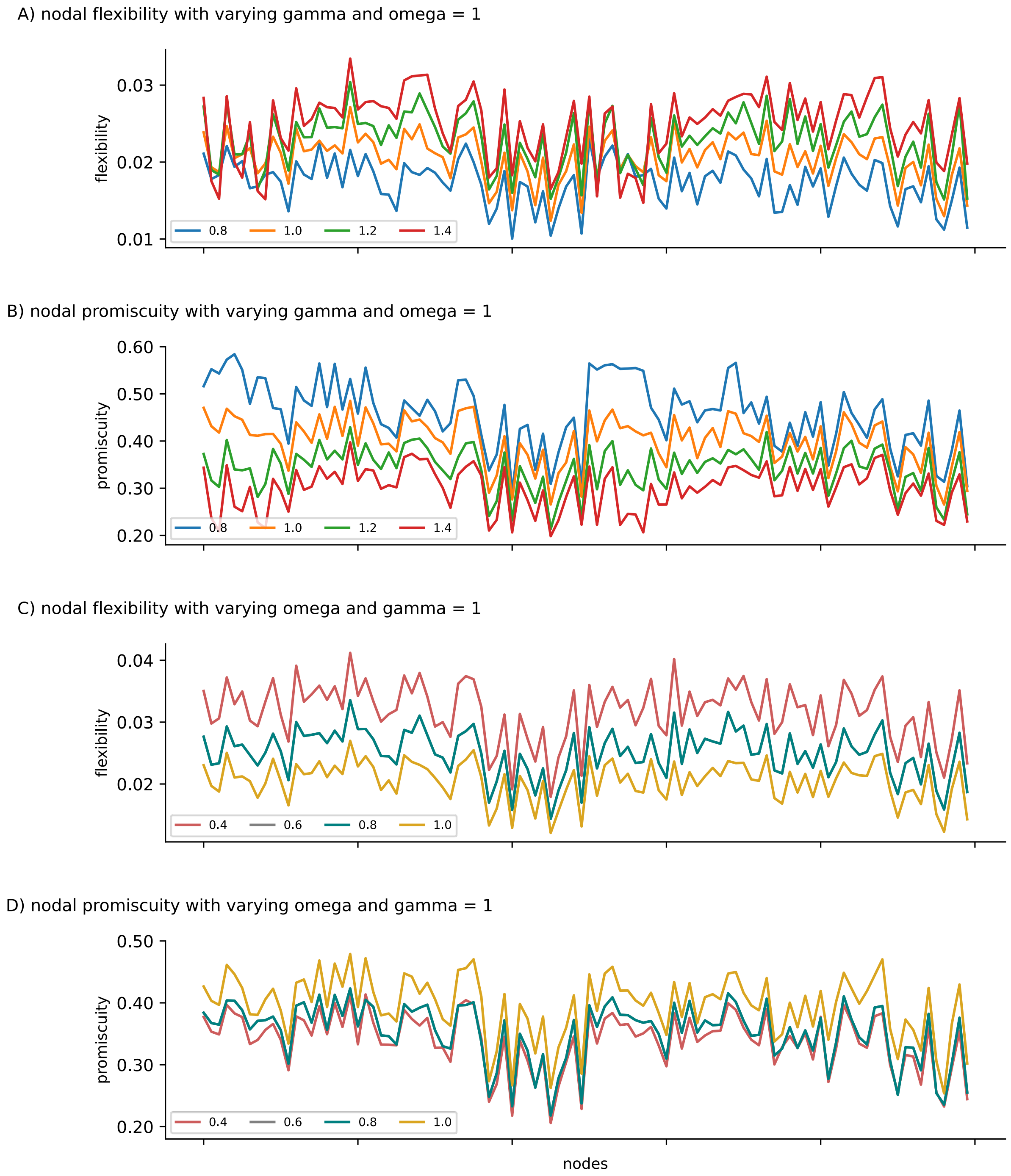


**Figure S3**. Influence of varying the intra- (gamma) and interlayer (omega) parameter while maximizing the multilayer modularity. A) Node flexibility and B) node promiscuity as a function of varying gamma while fixing omega = 1. C) Node flexibility and D) node promiscuity as a function of varying omega while fixing gamma = 1.

**Effect of different intra- and interlayer parameters on the correlation between measures of brain dynamics and psychological resilience**

While varying the interlayer parameter $\gamma$ from 0.8 to 1.4 in steps of 0.2 and computing nodal flexibility and promiscuity based on the original data (TR= .675s; Schaefer100 atlas), we did observe one significant correlation between flexibility of a visual node (LH_Vis_7) and the BRS ($r$ = .50, $p$ = .02) when using a parameter value of $\gamma$ = 1.4. For the remaining two levels of the intralayer parameter (note that we do not report $\gamma = 1$ here, as it refers to the initial analysis in the manuscript) we did not observe any significant correlations. The significant correlation indicates that higher flexibility is associated with higher self-reported resilience, which is directly opposed to what has been reported before (e.g., Long et al., 2019; Paban et al., 2019). Furthermore, across levels of $\gamma$, there were no significant correlations between node promiscuity and the resilience questionnaires (all $p$ > .35). We did not observe any significant correlation between node flexibility (all $p$ > .10) or node promiscuity (all $p$ > .11) and resilience while setting $\omega$ to either 0.4, 0.6, or 0.8.

### Supplementary References


Abraham, A., Pedregosa, F., Eickenberg, M., Gervais, P., Mueller, A., Kossaifi, J., Gramfort, A., Thirion, B., & Varoquaux, G. (2014). Machine learning for neuroimaging with scikit-learn. *Frontiers in Neuroinformatics*, *8*. https://doi.org/10.3389/fninf.2014.00014

Avants, B. B., Epstein, C. L., Grossman, M., & Gee, J. C. (2008). Symmetric diffeomorphic image registration with cross-correlation: Evaluating automated labeling of elderly and neurodegenerative brain. *Medical Image Analysis*, *12*(1), 26–41. https://doi.org/10.1016/j.media.2007.06.004

Behzadi, Y., Restom, K., Liau, J., & Liu, T. T. (2007). A component based noise correction method (CompCor) for BOLD and perfusion based fMRI. *NeuroImage*, *37*(1), 90–101. https://doi.org/10.1016/j.neuroimage.2007.04.042

Cox, R. W., & Hyde, J. S. (1997). Software tools for analysis and visualization of fMRI data. *NMR in Biomedicine*, *10*(4–5), 171–178. https://doi.org/10.1002/(SICI)1099-1492(199706/08)10:4/5<171::AID-NBM453>3.0.CO;2-L

Dale, A. M., Fischl, B., & Sereno, M. I. (1999). Cortical Surface-Based Analysis: I. Segmentation and Surface Reconstruction. *NeuroImage*, *9*(2), 179–194. https://doi.org/10.1006/nimg.1998.0395

Esteban, O., Blair, R., Markiewicz, C. J., Berleant, S. L., Moodie, C., Ma, F., Isik, A. I., Erramuzpe, A., Kent, M., James D. andGoncalves, DuPre, E., Sitek, K. R., Gomez, D. E. P., Lurie, D. J., Ye, Z., Poldrack, R. A., & Gorgolewski, K. J. (2018a). FMRIPrep. *Software*. https://doi.org/10.5281/zenodo.852659

Esteban, O., Markiewicz, C., Blair, R. W., Moodie, C., Isik, A. I., Erramuzpe Aliaga, A., Kent, J., Goncalves, M., DuPre, E., Snyder, M., Oya, H., Ghosh, S., Wright, J., Durnez, J., Poldrack, R., & Gorgolewski, K. J. (2018b). fMRIPrep: A robust preprocessing pipeline for functional MRI. *Nature Methods*. https://doi.org/10.1038/s41592-018-0235-4

Evans, A., Janke, A., Collins, D., & Baillet, S. (2012). Brain templates and atlases. *NeuroImage*, *62*(2), 911–922. https://doi.org/10.1016/j.neuroimage.2012.01.024

Fonov, V., Evans, A., McKinstry, R., Almli, C., & Collins, D. (2009). Unbiased nonlinear average age-appropriate brain templates from birth to adulthood. *NeuroImage*, *47, Supplement 1*, S102. https://doi.org/10.1016/S1053-8119(09)70884-5

Fortunato, S., & Barthelemy, M. (2007). Resolution limit in community detection. *Proceedings of the National Academy of Sciences*, *104*(1), 36–41. https://doi.org/10.1073/pnas.0605965104

Furtunato, S. (2010). Community detection in graphs. *Physics Reports*, *486*(3–5), 75–174. https://doi.org/10.1016/j.physrep.2009.11.002

Glasser, M. F., Sotiropoulos, S. N., Wilson, J. A., Coalson, T. S., Fischl, B., Andersson, J. L., Xu, J., Jbabdi, S., Webster, M., Polimeni, J. R., Van Essen, D. C., & Jenkinson, M. (2013). The minimal preprocessing pipelines for the Human Connectome Project. *NeuroImage*, *80*, 105–124. https://doi.org/10.1016/j.neuroimage.2013.04.127

Gorgolewski, K., Burns, C. D., Madison, C., Clark, D., Halchenko, Y. O., Waskom, M. L., & Ghosh, S. (2011). Nipype: A flexible, lightweight and extensible neuroimaging data processing framework in Python. *Frontiers in Neuroinformatics*, *5*, 13. https://doi.org/10.3389/fninf.2011.00013

Gorgolewski, K. J., Auer, T., Calhoun, V. D., Craddock, R. C., Das, S., Duff, E. P., Flandin, G., Ghosh, S. S., Glatard, T., Halchenko, Y. O., Handwerker, D. A., Hanke, M., Keator, D., Li, X., Michael, Z., Maumet, C., Nichols, B. N., Nichols, T. E., Pellman, J., … Poldrack, R. A. (2016). The brain imaging data structure, a format for organizing and describing outputs of neuroimaging experiments. *Scientific Data*, *3*(1), 160044. https://doi.org/10.1038/sdata.2016.44

Gorgolewski, K. J., Esteban, O., Markiewicz, C. J., Ziegler, E., Ellis, D. G., Notter, M. P., Jarecka, D., Johnson, H., Burns, C., Manhães-Savio, A., Hamalainen, C., Yvernault, B., Salo, T., Jordan, K., Goncalves, M., Waskom, M., Clark, D., Wong, J., Loney, F., … Ghosh, S. (2018). Nipype. *Software*. https://doi.org/10.5281/zenodo.596855

Greve, D. N., & Fischl, B. (2009). Accurate and robust brain image alignment using boundary-based registration. *NeuroImage*, *48*(1), 63–72. https://doi.org/10.1016/j.neuroimage.2009.06.060

Huntenburg, J. M. (2014). *Evaluating nonlinear coregistration of BOLD EPI and T1w images* [Master's Thesis, Freie Universität]. http://hdl.handle.net/11858/00-001M-0000-002B-1CB5-A

Jenkinson, M., Bannister, P., Brady, M., & Smith, S. (2002). Improved Optimization for the Robust and Accurate Linear Registration and Motion Correction of Brain Images. *NeuroImage*, *17*(2), 825–841. https://doi.org/10.1006/nimg.2002.1132

Jenkinson, M., & Smith, S. (2001). A global optimisation method for robust affine registration of brain images. *Medical Image Analysis*, *5*(2), 143–156. https://doi.org/10.1016/S1361-8415(01)00036-6

Klein, A., Ghosh, S. S., Bao, F. S., Giard, J., Häme, Y., Stavsky, E., Lee, N., Rossa, B., Reuter, M., Neto, E. C., & Keshavan, A. (2017). Mindboggling morphometry of human brains. *PLOS Computational Biology*, *13*(2), e1005350. https://doi.org/10.1371/journal.pcbi.1005350

Lanczos, C. (1964). Evaluation of Noisy Data. *Journal of the Society for Industrial and Applied Mathematics Series B Numerical Analysis*, *1*(1), 76–85. https://doi.org/10.1137/0701007

Long, Y., Chen, C., Deng, M., Huang, X., Tan, W., Zhang, L., Fan, Z., & Liu, Z. (2019). Psychological resilience negatively correlates with resting-state brain network flexibility in young healthy adults: A dynamic functional magnetic resonance imaging study. *Annals of Translational Medicine*, *7*(24), 809. https://doi.org/10.21037/atm.2019.12.45

Markiewicz, C. J., Goncalves, M., Blair, R. W., Gorgolewski, K. J., Poldrack, R. A., & Esteban, O. (2020). SDCflows: Susceptibility Distortion Correction workFLOWS. Zenodo. https://doi.org/10.5281/zenodo.3668335

Mazziotta, J. C., Toga, A. W., Evans, A., Fox, P., & Lancaster, J. (1995). A Probabilistic Atlas of the Human Brain: Theory and Rationale for Its Development: The International Consortium for Brain Mapping (ICBM). *NeuroImage*, *2*(2, Part A), 89–101. https://doi.org/10.1006/nimg.1995.1012

Mucha, P. J., Richardson, T., Macon, K., Porter, M. A., & Onnela, J.-P. (2010). Community Structure in Time-Dependent, Multiscale, and Multiplex Networks. *Science*, *328*(5980), 876–878. https://doi.org/10.1126/science.1184819

Paban, V., Modolo, J., Mheich, A., & Hassan, M. (2019). Psychological resilience correlates with EEG source-space brain network flexibility. *Network Neuroscience*, *3*(2), 539–550. https://doi.org/10.1162/netn_a_00079

Pedersen, M., Zalesky, A., Omidvarnia, A., & Jackson, G. D. (2018). Multilayer network switching rate predicts brain performance. *Proceedings of the National Academy of Sciences*, *115*(52), 13376–13381. https://doi.org/10.1073/pnas.1814785115

Posse, S., Wiese, S., Gembris, D., Mathiak, K., Kessler, C., Grosse-Ruyken, M.-L., Elghahwagi, B., Richards, T., Dager, S. R., & Kiselev, V. G. (1999). Enhancement of BOLD-contrast sensitivity by single-shot multi-echo functional MR imaging. *Magnetic Resonance in Medicine*, *42*(1), 87–97. https://doi.org/10.1002/(SICI)1522-2594(199907)42:1<87::AID-MRM13>3.0.CO;2-O

Power, J. D., Cohen, A. L., Nelson, S. M., Wig, G. S., Barnes, K. A., Church, J. A., Vogel, A. C., Laumann, T. O., Miezin, F. M., Schlaggar, B. L., & Petersen, S. E. (2011). Functional network organization of the human brain. *Neuron*, *72*(4),

665–678. https://doi.org/10.1016/j.neuron.2011.09.006

Power, J. D., Mitra, A., Laumann, T. O., Snyder, A. Z., Schlaggar, B. L., & Petersen, S. E. (2014). Methods to detect, characterize, and remove motion artifact in resting state fMRI. *NeuroImage*, *84*(Supplement C), 320–341. https://doi.org/10.1016/j.neuroimage.2013.08.048

Pruim, R. H. R., Mennes, M., van Rooij, D., Llera, A., Buitelaar, J. K., & Beckmann, C. F. (2015). ICA-AROMA: A robust ICA-based strategy for removing motion artifacts from fMRI data. *NeuroImage*, *112*(Supplement C), 267–277. https://doi.org/10.1016/j.neuroimage.2015.02.064

Reuter, M., Rosas, H. D., & Fischl, B. (2010). Highly accurate inverse consistent registration: A robust approach. *NeuroImage*, *53*(4), 1181–1196. https://doi.org/10.1016/j.neuroimage.2010.07.020

Satterthwaite, T. D., Elliott, M. A., Gerraty, R. T., Ruparel, K., Loughead, J., Calkins, M. E., Eickhoff, S. B., Hakonarson, H., Gur, R. C., Gur, R. E., & Wolf, D. H. (2013). An improved framework for confound regression and filtering for control of motion artifact in the preprocessing of resting-state functional connectivity data. *NeuroImage*, *64*(1), 240–256. https://doi.org/10.1016/j.neuroimage.2012.08.052

Treiber, J. M., White, N. S., Steed, T. C., Bartsch, H., Holland, D., Farid, N., McDonald, C. R., Carter, B. S., Dale, A. M., & Chen, C. C. (2016). Characterization and Correction of Geometric Distortions in 814 Diffusion Weighted Images. *PLOS ONE*, *11*(3), e0152472. https://doi.org/10.1371/journal.pone.0152472

Tustison, N. J., Avants, B. B., Cook, P. A., Zheng, Y., Egan, A., Yushkevich, P. A., & Gee, J. C. (2010). N4ITK: Improved N3 Bias Correction. *IEEE Transactions on Medical Imaging*, *29*(6), 1310–1320. https://doi.org/10.1109/TMI.2010.2046908

Tzourio-Mazoyer, N., Landeau, B., Papathanassiou, D., Crivello, F., Etard, O., Delcroix, N., Mazoyer, B., & Joliot, M. (2002). Automated anatomical labeling of activations in SPM using a macroscopic anatomical parcellation of the MNI MRI single-subject brain. *NeuroImage*, *15*(1), 273–289. https://doi.org/10.1006/nimg.2001.0978

Vaiana, M., & Muldoon, S. (2018). Resolution Limits for Detecting Community Changes in Multilayer Networks. *arXiv:1803.03597 [physics]*.

http://arxiv.org/abs/1803.03597
Wang, S., Peterson, D. J., Gatenby, J. C., Li, W., Grabowski, T. J., & Madhyastha, T. M. (2017). Evaluation of Field Map and Nonlinear Registration Methods for Correction of Susceptibility Artifacts in Diffusion MRI. *Frontiers in Neuroinformatics*, *11*. https://doi.org/10.3389/fninf.2017.00017
Yin, W., Li, T., Hung, S.-C., Zhang, H., Wang, L., Shen, D., Zhu, H., Mucha, P. J., Cohen, J. R., & Lin, W. (2020). The emergence of a functionally flexible brain during early infancy. *Proceedings of the National Academy of Sciences*, 202002645. https://doi.org/10.1073/pnas.2002645117
Zhang, Y., Brady, M., & Smith, S. (2001). Segmentation of brain MR images through a hidden Markov random field model and the expectation-maximization algorithm. *IEEE Transactions on Medical Imaging*, *20*(1), 45–57. https://doi.org/10.1109/42.906424
